# White matter–cognition mapping in pediatric cancer survivors: a two-cohort NODDI-Bingham study

**DOI:** 10.64898/2026.09.25.754456

**Authors:** Shounak Nandi, Faye McKenna, Sonya Henry, Siddharth Nayak, Timothy Duong

## Abstract

Childhood cancer survivors (CS) treated with chemotherapy frequently experience long-term cognitive impairments, commonly referred to as CRCI. Whether these deficits reflect disrupted white matter microstructure, altered microstructure–cognition relationships, or both remains unclear. Here, we applied the NODDI-Bingham model, which quantifies neurite density (NDI), isotropic free water (ISO), and orientation dispersion along primary (ODIp), secondary (ODIs), and total (ODIt) fiber axes, to characterize biophysically specific white matter microstructure in CS and examine its relationship to cognition.

Twenty-one pediatric cancer survivors and 21 age-matched healthy controls (HC) underwent multi-shell diffusion MRI. Voxel-wise NODDI-Bingham analyses were performed within JHU atlas regions. Group differences were tested by ANCOVA, and microstructure–cognition associations were examined with the NIH Toolbox Crystallized, Fluid, and Early Childhood composites using group-specific regressions. Confirmatory analyses were performed in an age-matched subsample of the HCP (n = 679).

Notably, CS showed reduced NDI in the corona radiata, cerebral peduncles, and corpus callosum, with elevated ODIt and ODIp in the middle cerebellar peduncle, pathways that subserve processing speed and interhemispheric integration, cognitive domains among the most vulnerable in CRCI. Group-specific regressions revealed qualitatively different microstructure–cognition relationships: HC exhibited mixed positive and negative relationships across four of five metrics (138 clusters spanning approximately 305,000 voxels; peak Cohen’s d up to 3.00), whereas CS showed a predominantly uniformly negative pattern (67 clusters across 57,000 voxels), with no positive clusters for ISO, ODIt, ODIp, or ODIs, thus highlighting that reduced dispersion and extracellular free water are linked to better cognitive performance in CS. HCP analyses reproduced the HC pattern across all five metrics at proportionally smaller effect sizes, ruling out sample-size effects.

NODDI-Bingham identifies two aspects of chemotherapy-related white matter pathology that standard diffusion metrics cannot resolve. First, pediatric cancer survivors show selective elevation of ODIp and ODIt in the cerebellar peduncles and reduced NDI in the corona radiata and corpus callosum, localizing group-level injury to specific fanning and neurite density signatures. Second, whereas in healthy controls white matter shows mixed positive and negative microstructure–cognition relationships, survivors retain only negative relationships for ODIp and ODIt and a diminished mixed pattern for NDI, such that in the chemotherapy-exposed brain, only reduced neurite density and increased primary-axis fanning relate to cognitive impairment. These findings implicate a biophysically specific restructuring of white matter–cognition relationships as a candidate mechanism for CRCI and support NODDI-Bingham as a targeted biomarker panel for monitoring and intervention trials.

**Abbreviated summary:** Nandi, McKenna et al. apply the NODDI-Bingham diffusion model to pediatric cancer survivors and controls, showing chemotherapy is linked to selective white matter changes and a restructuring of microstructure–cognition relationships. The findings position NODDI-Bingham as a candidate biomarker panel for tracking and treating cancer-related cognitive impairment.

## Introduction

Advances in pediatric oncology have yielded five-year survival rates exceeding 85% for many childhood cancers, yet long-term neurocognitive morbidity remains a significant concern^1^. Survivors of childhood cancer, particularly those treated with systemic chemotherapy, frequently demonstrate persistent deficits in processing speed, working memory, attention, and executive function that can emerge years after treatment cessation and persist into adulthood^2,3^. These impairments, collectively referred to as chemotherapy-related cognitive impairment (CRCI), have substantial consequences for educational attainment, vocational functioning, and quality of life^4^. Understanding the neural substrates of CRCI is therefore critical for developing targeted interventions and monitoring strategies for this growing survivor population.

White matter (WM) integrity loss is a primary effect of chemotherapy-induced neurotoxicity^5^. Preclinical evidence demonstrates that chemotherapeutic agents, particularly antimetabolites such as methotrexate, directly damage oligodendrocyte precursor cells, reduce their capacity to differentiate into mature myelinating oligodendrocytes, and trigger neuroinflammatory cascades that further compromise myelin homeostasis^6,7^. Clinically, diffusion tensor imaging (DTI) studies have consistently reported reduced fractional anisotropy (FA) and increased mean diffusivity in major WM tracts of cancer survivors, including the corpus callosum, corona radiata, and frontoparietal pathways, with these microstructural alterations correlating with deficits in processing speed, attention, and executive function^5,8–12^. In pediatric cancer populations specifically, DTI studies have documented reduced FA in cerebellar, brainstem, and long-range commissural pathways^10,12,13^. Our own prior work using free-water-corrected intravoxel incoherent motion diffusion imaging (IVIM-FWI) in a partially overlapping pediatric cancer survivor cohort detected widespread group differences in perfusion fraction, free water fraction, and tissue anisotropy, including in the cerebellar peduncle, corona radiata, and corpus callosum^14^. However, conventional DTI metrics and their free-water corrected refinements are inherently non-specific: changes in FA may reflect alterations in axonal density, myelination, fiber coherence, or extracellular water content, making it difficult to identify the precise microstructural substrates of cognitive impairment^15^.

Neurite orientation dispersion and density imaging (NODDI) addresses this limitation by fitting a multi-compartment biophysical model to multi-shell diffusion data, separately estimating neurite density (NDI), orientation dispersion (ODI), and isotropic free water volume fraction (ISO)^16^. The Bingham extension of NODDI (NODDI-Bingham) further decomposes orientation dispersion into primary (ODIp) and secondary (ODIs) components, along with total dispersion (ODIt)^17^ (Fig. 1). This decomposition is neurobiologically meaningful: ODIp captures the spread of neurites along the primary dispersion axis (the plane in which fibers fan or bend most prominently), and is sensitive to the coherence of fiber arrangements that depend on intact myelin sheaths and organized axonal packing^18^. ODIs reflect dispersion perpendicular to this plane, capturing a geometrically independent aspect of fiber organization^18^. Together, the five NODDI-Bingham metrics (NDI, ISO, ODIt, ODIp, and ODIs) provide a panel of directionally specific microstructural markers, each potentially sensitive to a different aspect of chemotherapy-related white matter pathology. This biological specificity also creates an opportunity beyond group-level comparisons: because each metric indexes a distinct aspect of tissue organization, examining how each metric relates to cognition within a group offers a window onto the *architecture* of microstructure–cognition coupling, and onto how that architecture may be altered by disease.

**Figure 1.**
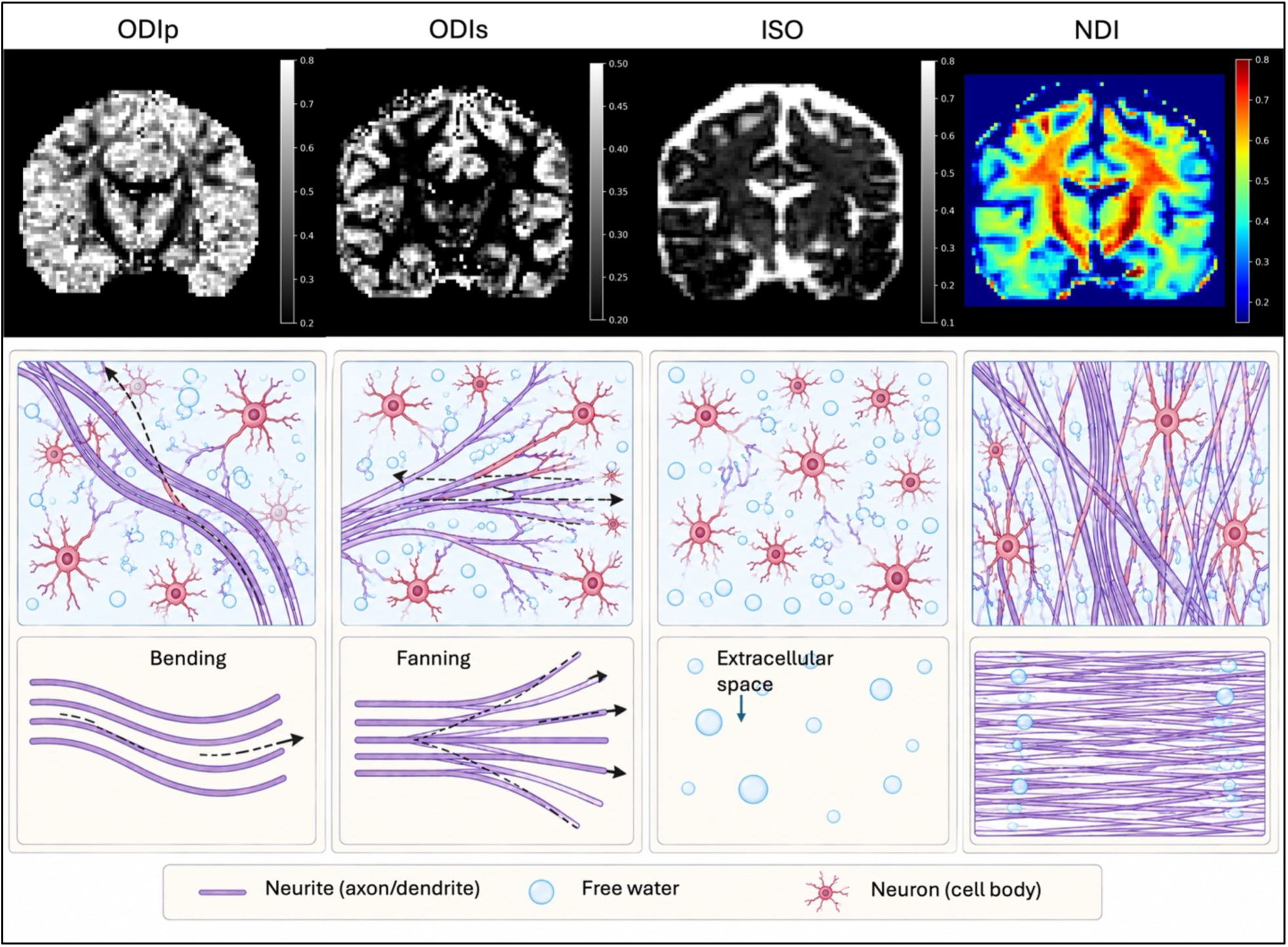
Schematic representation of the Bingham-NODDI metrics showing ODIp, ODIs, ISO, and NDI. Top row shows the diffusion maps of the ODIp, ODIs, ISO and NDI. The ODIp shows the primary bending, visually seen in the corpus callosum, while the secondary fanning is captured by ODIs towards the cortical areas highlighted by red arrowheads. ISO shows the free water in the brain, primarily highlighted around CSF, and NDI shows the neurite density which are of greater magnitude along the white matter structures (shade of red). Bottom row illustrates these metrics schematically. Note, the dispersion along the primary and secondary axes are usually anisotropic, where the ODIp follows a dispersion along the bend of neurites, principally along the primary axis, while the ODIs follows a dispersion along the secondary axis. The Bingham distribution is shown in the bottom row, which determines the direction of the dispersion of the orientation distribution function. Colorscale 0.8 matched across three metrics.

Two gaps in the existing literature motivate the present study. First, the only prior application of NODDI to pediatric cancer survivors was by Sleurs et al. (2018), who reported elevated isotropic volume fraction (VISO) distributed throughout central white matter and focally higher NDI in the corticospinal tract of childhood bone and soft tissue sarcoma survivors treated with systemic intravenous chemotherapy, with no significant group differences in ODI^19^. However, that analysis used the standard Watson distribution, which is optimized for gray matter and assumes symmetric dispersion and cannot distinguish between primary and secondary dispersion patterns^17^. No study has applied the Bingham extension of NODDI to pediatric cancer survivors, leaving the directional specificity of orientation dispersion changes (and their relationship to cognition) uncharacterized in this population. Second, and more fundamentally, prior neuroimaging work on chemotherapy-related white matter pathology has focused almost exclusively on mean group differences, not on how the relationship between microstructure and cognition is organized within each group^20^. Whether chemotherapy’s cognitive impact reflects disrupted microstructure, disrupted microstructure–cognition relationships, or both, therefore remains unclear. Addressing this requires a framework that can characterize how multiple microstructural metrics relate to cognitive performance within each group separately, and an external normative cohort against which the control-group patterns can be validated.

The present study addresses these gaps by applying NODDI-Bingham to pediatric cancer survivors (CS) and age-matched controls (HC), with confirmatory analyses in an age-matched subsample (*n* = 679) of the Human Connectome Project (HCP) 1200 dataset^21^. Our central question was whether chemotherapy disrupts the architecture of microstructure–cognition coupling, not only the mean levels of microstructural indices. Three aims followed from this question: (1) to characterize group differences in NODDI-Bingham metrics between CS and HC; (2) to examine whether the pattern of microstructure–cognition associations differs between groups; and (3) to test whether the healthy control pattern replicates in the HCP normative sample, providing external validation that any CS-specific disruption is not a small-sample artifact. We hypothesized that cancer survivors would show reduced neurite density and increased isotropic volume fraction and orientation dispersion in cerebellar and cerebral white matter pathways, and that the microstructure–cognition associations observed in healthy controls would be specifically disrupted in the CS group.

## Materials and Methods

### Participant Recruitment

The study was approved by the Albert Einstein College of Medicine institutional review board (IRB-2020-12265), and all participants and/or their legal guardians provided written informed consent. Ethical principles were strictly adhered to, including maintaining participant anonymity and minimizing potential risks associated with the research.

Participants (15-35 years old) included pediatric cancer patients and age-matched healthy controls. Cancer patients were childhood cancer survivors who were 10-25 years post-chemotherapy. Eligible cancer types included acute lymphoblastic leukemia (ALL), acute myeloid leukemia, neuroblastoma, non-CNS solid tumors, and Hodgkin lymphoma. See Supplementary Methods for details.

### NIH Toolbox: Cognition Battery

Cognitive performance was assessed using the NIH Toolbox Cognition Battery^22^, which included the Fluid Intelligence and Early Childhood composite scores, as well as the Flanker Inhibitory Control and Attention test and the Pattern Comparison Processing Speed test. See Supplementary Methods for details.

### Magnetic Resonance Imaging

MRI data were acquired on a 3T MRI and a 32-element head-neck coil. Three-dimensional T1-weighted images (T1w) were acquired with a voxel size of 0.87 mm^3^ and used to align and register with diffusion MRI (dMRI) images. Diffusion MRI data were acquired with a voxel size of 2 mm^3^ and b values = 0 (6), 300 (6), 800 (27), and 2000 (30) s/mm^2^. See Supplementary Methods for details.

Diffusion images followed standard preprocessing steps (see Supplementary Methods). The multi-compartment NODDI-Bingham model proposed by Tariq et al. (2016) was implemented using Diffusion Microstructure Imaging in Python (DMIPY)^17,23^. Unlike the standard NODDI model that uses the Watson distribution to compute the metrics, the NODDI-Bingham uses the Bingham distribution model to account for the fanning and bending of the fibers that are usually common and account for anisotropy. Thus using this model, additional metrics are deduced: Neurite Density Index (NDI), Isotropic volume fraction (ISO), Orientation dispersion Indices (ODIt, ODIp, and ODIs):

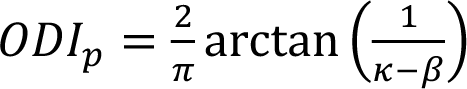 along axis **μ**2, 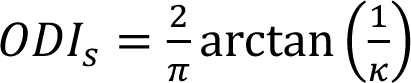 along axis **μ**3, where **μ**2, and **µ**3 are principal and secondary dispersion orientation, respectively. *β* denotes the ratio of primary to secondary dispersion, a value of 0 means isotropic dispersion (recovering Watson/NODDI), while higher values indicate more asymmetric fanning, and *κ* denotes the concentration parameter^17^.

The overall dispersion index, ODIt is computed as:

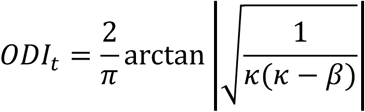

The FreeSurfer software (version 7.4)^24^ was used to parcellate the WM ROIs of the Johns Hopkins University (JHU) atlas^25^ and warp dMRI maps into the common MNI space.

### Statistical Analyses

Group differences in clinical cognitive measures were assessed using independent-samples *t*-tests, with a significance threshold of *P* < 0.05, in Python version 3.9. Voxel-wise analyses were conducted on the dMRI maps in MNI space, masking for the *a priori* ROIs using FSL randomise with 500 permutations and a *P* < 0.05 cluster-forming Family-wise error (FWE)-corrected threshold^26^. First, analyses of covariance (ANCOVA) controlling for age and sex were conducted to compare the CS and HC groups. Subsequently, voxel-wise regression analyses were performed in each group to assess potential associations between NODDI-Bingham metrics and cognitive test scores, controlling for age and sex. Effect sizes were calculated using Cohen’s *d*^27^. Voxel-wise regression analyses were also performed in the whole group, additionally controlling for group membership (see Supplementary Materials).

#### HCP Confirmatory Analyses

To evaluate whether the microstructure–cognition associations observed in the HC group generalized to a large independent normative sample, we conducted confirmatory voxel-wise regression analyses in the HCP 1200 dataset on an age-matched subsample restricted to participants aged 22–30 years (*n* = 679; Table 2)^21^. NODDI-Bingham maps were computed for HCP participants with available multi-shell diffusion MRI data (b = 0, 1000, 2000, 3000 s/mm²) using the same DMIPY pipeline applied to the primary cohort. Regressions were performed between each NODDI-Bingham metric and the same three NIH Toolbox cognitive composites (CCC, Fluid Composite, Early Childhood Composite) using FSL randomise with 500 permutations and *P* < .05 FWE-corrected cluster-forming threshold, controlling for age and sex. One participant’s data were excluded due to missing b-values and b-vectors.

## Results

Demographic and clinical characteristics for both groups are presented in Table 1. The chemotherapy (CS; *n* = 21) and healthy control (HC; *n* = 21) groups did not differ significantly in age (*P* = 0.246), though they differed in racial and ethnic distribution (both *P* = 0.003). Cancer survivors demonstrated significantly lower scores on the Cognitive Crystallized Composite (CCC; *P* < 0.001, *d* = 1.31), Fluid Composite (*P* = 0.006, *d* = 0.90), and Cognition Total Composite (*P* < 0.001, *d* = 1.22) relative to controls. Among the individual subtests, CS scored significantly lower on the Pattern Comparison Processing Speed test (*P* = 0.001, *d* = 1.23), Oral Reading Recognition test (*P* < 0.001, *d* = 1.49), List Sorting Working Memory test (*P* = 0.010, *d* = 0.89), Flanker Inhibitory Control and Attention test (*P* = 0.025, *d* = 0.79), and Picture Vocabulary test (*P* = 0.033, *d* = 0.76). The Early Childhood Composite trended lower in CS but did not reach significance (*P* = 0.068, *d* = 0.58), while the Dimensional Change Card Sort and Picture Sequence tests did not differ between groups (*P* = 0.336 and *P* = 0.540, respectively).

**Table 1.**
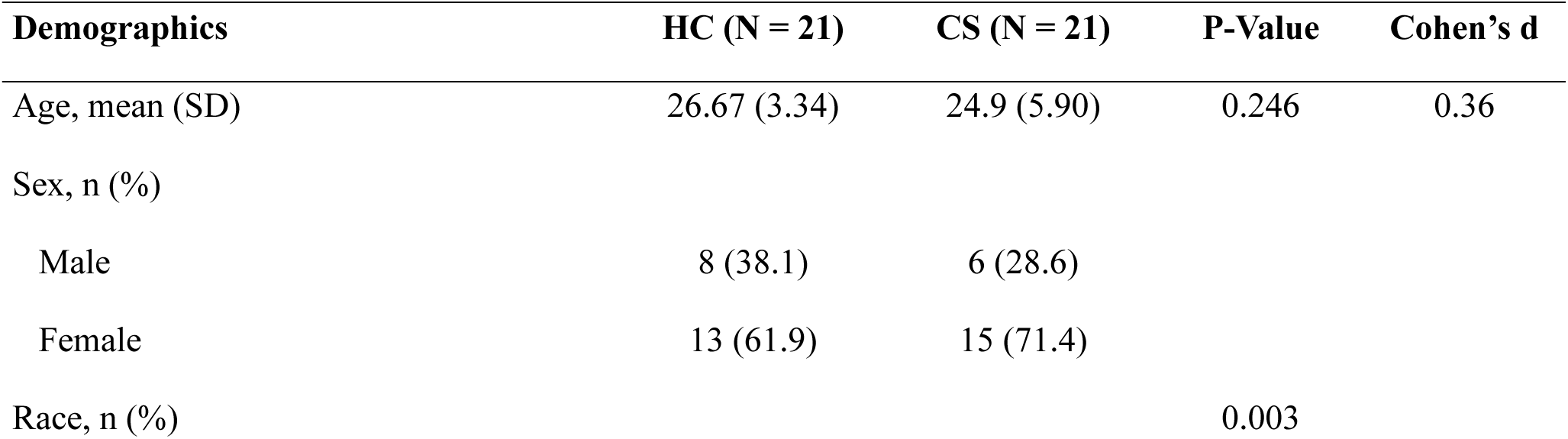
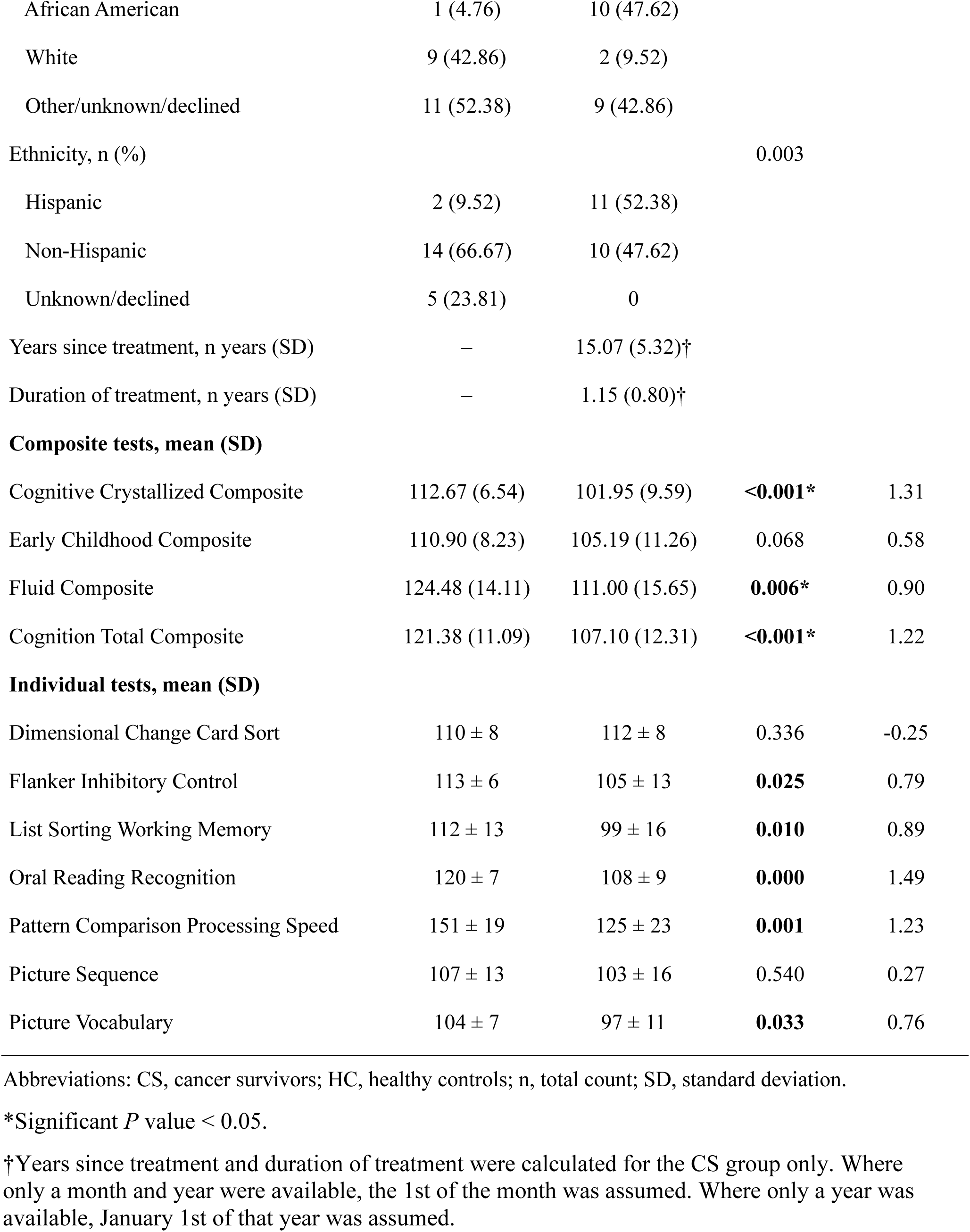
Demographic and clinical characteristics of the control and chemo groups.

| Demographics | HC (N = 21) | CS (N = 21) | P-Value | Cohen's d |
| --- | --- | --- | --- | --- |
| Age, mean (SD) | 26.67 (3.34) | 24.9 (5.90) | 0.246 | 0.36 |
| Sex, n (%) |  |  |  |  |
| Male | 8 (38.1) | 6 (28.6) |  |  |
| Female | 13 (61.9) | 15 (71.4) |  |  |
| Race, n (%) |  |  | 0.003 |  |
| African American | 1 (4.76) | 10 (47.62) |  |  |
| White | 9 (42.86) | 2 (9.52) |  |  |
| Other/unknown/declined | 11 (52.38) | 9 (42.86) |  |  |
| Ethnicity, n (%) |  |  | 0.003 |  |
| Hispanic | 2 (9.52) | 11 (52.38) |  |  |
| Non-Hispanic | 14 (66.67) | 10 (47.62) |  |  |
| Unknown/declined | 5 (23.81) | 0 |  |  |
| Years since treatment, n years (SD) | – | 15.07 (5.32)† |  |  |
| Duration of treatment, n years (SD) | – | 1.15 (0.80)† |  |  |
| **Composite tests, mean (SD)** |  |  |  |  |
| Cognitive Crystallized Composite | 112.67 (6.54) | 101.95 (9.59) | <0.001* | 1.31 |
| Early Childhood Composite | 110.90 (8.23) | 105.19 (11.26) | 0.068 | 0.58 |
| Fluid Composite | 124.48 (14.11) | 111.00 (15.65) | 0.006* | 0.90 |
| Cognition Total Composite | 121.38 (11.09) | 107.10 (12.31) | <0.001* | 1.22 |
| **Individual tests, mean (SD)** |  |  |  |  |
| Dimensional Change Card Sort | 110 ± 8 | 112 ± 8 | 0.336 | -0.25 |
| Flanker Inhibitory Control | 113 ± 6 | 105 ± 13 | 0.025 | 0.79 |
| List Sorting Working Memory | 112 ± 13 | 99 ± 16 | 0.010 | 0.89 |
| Oral Reading Recognition | 120 ± 7 | 108 ± 9 | 0.000 | 1.49 |
| Pattern Comparison Processing Speed | 151 ± 19 | 125 ± 23 | 0.001 | 1.23 |
| Picture Sequence | 107 ± 13 | 103 ± 16 | 0.540 | 0.27 |
| Picture Vocabulary | 104 ± 7 | 97 ± 11 | 0.033 | 0.76 |

Abbreviations: CS, cancer survivors; HC, healthy controls; n, total count; SD, standard deviation.
\*Significant *P* value < 0.05.
†Years since treatment and duration of treatment were calculated for the CS group only. Where only a month and year were available, the 1st of the month was assumed. Where only a year was available, January 1st of that year was assumed.

Demographic and clinical characteristics differences for the HCP group with the CS and HC groups are tabulated in Table 2. The HCP group controls (*n* = 679) showed significantly greater CCC scores (*P* = 0.048, *d* = 0.44) but did not show significance in Early Childhood Composite (*P* = 0.171, *d* = 0.39) and Fluid Composite scores (*P* = 0.861, *d* = 0.06) as compared to the CS group. The HC group, however, showed higher scores than the HCP group in all three composite scores, CCC (*P* < 0.001, *d* = 0.73), Early Childhood Composite (*P* = 0.677, *d* = 0.09), and Fluid Composite (*P* = 0.003, *d* = 0.75).

**Table 2.** Comparison of the cognitive scores of the HCP Age-matched controls (HCP) versus the CS group and HC group.

| Measure | HCP_sub<br>(n = 679) | CS (n =<br>21) | HC | P-value,<br>HCP vs<br>CS | Cohen's<br>d | P-value,<br>HCP vs<br>HC | Cohen's<br>d |
| --- | --- | --- | --- | --- | --- | --- | --- |
| Age in years, mean (SD) | 26.54 (2.45) | 24.9 | 26.67 | - | - | - | - |
| Sex |  |  |  |  |  |  |  |
| *Male* | 351 (51.7%) | 6 | 8 | - | - | - | - |
| *Female* | 328 (48.3%) | 15 | 13 | - | - | - | - |
| Cognitive Crystallized<br>Composite, mean (SD) | 110.98<br>(17.26) | 103.42<br>(16.23) | **123.57**<br>**(13.57)** | **0.048** | 0.44 | **<0.001** | -0.73 |
| Early Childhood<br>Composite, mean (SD) | 107.14<br>(16.13) | 100.81<br>(20.27) | 108.61<br>(15.71) | 0.171 | 0.39 | 0.677 | -0.09 |
| Fluid Composite, mean<br>(SD) | 105.48<br>(17.34) | 104.52<br>(24.63) | 118.42<br>(17.42) | 0.861 | 0.06 | **0.003** | -0.75 |

Abbreviations: CCC, Cognitive Crystallized Composite; CS, cancer survivors; d, Cohen's d; HC, healthy controls; HCP, Human Connectome Project; HCP\_sub, age-matched HCP subsample; SD, standard deviation.

### Group Differences in NODDI-Bingham Metrics

Voxel-wise group comparisons revealed significant differences between CS and HC in multiple NODDI-Bingham metrics, predominantly localized to cerebellar and cerebral white matter pathways (Table 3). Whole-sample associations between NODDI-Bingham metrics and cognitive performance are reported in Supplementary Table 1.

**Table 3.** Significant voxel cluster group differences between HC and CS groups with contrast HC>CS in NODDI-Bingham metrics at the *P* < 0.05 FWE-corrected level.

| NODDI<br>metric | Region | Subregion | Peak t-stat | P-Value | Peak Cohen's<br>d | Voxels | Coordinates [X Y<br>Z] |
| --- | --- | --- | --- | --- | --- | --- | --- |
| **ODIp** | Corona radiata | Right<br>Posterior | -3.73 | <0.001 | 1.15 | 297 | [67 86 101] |
|  | Cerebellar<br>peduncle | Middle | -4.7 | <0.001 | 1.45 | 231 | [83 108 44] |
| **ODIt** | Cerebellar<br>peduncle | Middle | -3.97 | <0.001 | 1.23 | 552 | [76 89 40] |
|  |  | Middle | -4.41 | <0.001 | 1.36 | 314 | [83 108 44] |
|  |  | Middle | -3.86 | 0.003 | 1.19 | 252 | [76 103 40] |
|  |  | Middle | -3.38 | <0.001 | 1.04 | 492 | [81 80 44] |
|  |  | Middle | -4.6 | 0.004 | 1.42 | 240 | [83 108 44] |
|  | Cerebral peduncle | Right | -4.08 | <0.001 | 1.26 | 297 | [80 108 59] |
|  |  | Right | -4.11 | <0.001 | 1.27 | 326 | [80 109 59] |
| **NDI** | Corona radiata | Right Superior | 3.2 | <0.001 | 0.99 | 1222 | [71 122 112] |
|  | Cerebral peduncle | Right | 4.08 | <0.001 | 1.26 | 684 | [80 105 57] |
|  |  | Right | 4.54 | <0.001 | 1.4 | 907 | [80 105 57] |
|  | CC | Body | 3 | 0.002 | 0.93 | 541 | [105 112 110] |

| NODDI metric | Region | Subregion | Peak t-stat | P-Value | Peak Cohen's d | Voxels | Coordinates [X Y Z] |
| --- | --- | --- | --- | --- | --- | --- | --- |
|  | CC | Body | 3.06 | <0.001 | 0.94 | 709 | [71 104 107] |
|  | CC | Body | 3.02 | 0.011 | 0.93 | 437 | [105 112 110] |
|  | Corona radiata | Right Posterior | 3.01 | 0.005 | 0.93 | 498 | [101 93 99] |

White cells indicate a negative t-stat and grey cells indicate a positive t-stat. Significance was determined using a permutation cluster-wise $P < 0.050$ familywise error (FWE)-corrected threshold. Positive t statistic values indicate greater levels of diffusion signal in the healthy control (HC) group ( $n = 21$ ) compared to the CS group ( $n = 21$ ). *Abbreviations: [X Y Z], coordinates of the voxels on the MNI 1 mm brain atlas; CC, corpus callosum; CS, cancer survivors; FWE, familywise error; HC, healthy controls; ISO, isotropic volume fraction; NDI, neurite density index; ODIp, primary orientation dispersion index; ODIs, secondary orientation dispersion index; ODIt, total orientation dispersion index; t, t-statistic; WM, white matter.*

The CS group exhibited significantly greater total ODIt compared to HC in several clusters spanning the middle cerebellar peduncle and the cerebral peduncle (peak *t* = 4.60, *P* = 0.004, *d* = 1.42; 7 clusters, 2,473 total voxels). The primary orientation dispersion index (ODIp) showed a more complex pattern, with CS demonstrating lower ODIp than HC in the right posterior corona radiata (*t* = −3.73, *P* < 0.001, *d* = 1.15) and middle cerebellar peduncle (*t* = −4.70, *P* < 0.001, *d* = 1.45) (Table 3, Fig. 2).

**Figure 2.**
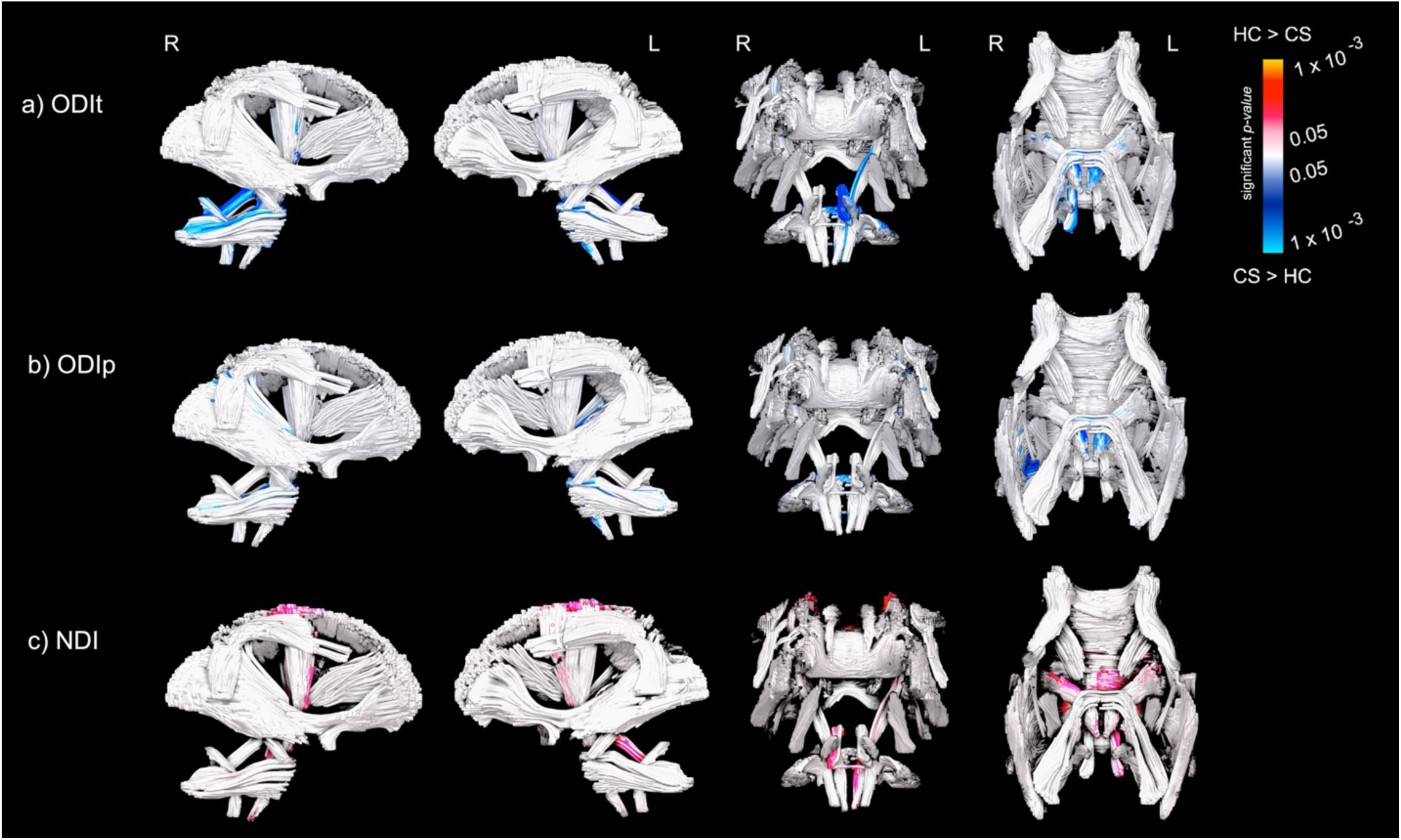
Significant voxelwise group differences in NODDI-Bingham white matter metrics between chemotherapy survivors (CS; n = 21) and healthy controls (HC; n = 21), covaried for age and sex. Analyses of covariance (ANCOVA) were conducted within white matter tracts defined by the JHU atlas, showing the main significance areas (a) total orientation dispersion index (ODIt), (b) primary orientation dispersion index (ODIp), and (c) neurite density index (NDI). Color indicates significant clusters at P < 0.05 familywise error (FWE), corrected threshold: red indicates higher values in HC relative to CS (HC > CS), and blue indicates lower values in HC relative to CS (CS > HC).

In contrast to the dispersion metrics, the NDI was significantly reduced in CS relative to HC across multiple white matter structures, including the right superior corona radiata (*t* = 3.20, *P* < 0.001, *d* = 0.99; 1,222 voxels), right cerebral peduncle (peak *t* = 4.54, *P* < 0.001, *d* = 1.40), and the body of the corpus callosum (CC; peak *t* = 3.06, *P* < 0.001, *d* = 0.94; 7 clusters, 4,997 total voxels). The regions showing group-level microstructural differences, the middle cerebellar peduncle, the cerebral peduncles, the corpus callosum, and the corona radiata, are the same regions in which altered microstructure–cognition relationships emerged in the group-specific regression analyses reported below (Table 3, Fig. 2).

### Group-Specific Associations Between NODDI-Bingham Metrics and Cognitive Performance

To examine whether brain–cognition relationships differed between groups, voxel-wise regression analyses were conducted separately within each group, controlling for age and sex (Tables 4 and 5, Fig. 3). The central finding of these analyses is that chemotherapy survivors showed not merely weaker but qualitatively different microstructure–cognition relationships than controls: the mixed positive and negative relationship that characterized healthy white matter, a pattern that was preserved when the HC regressions were replicated in the HCP cohort, was replaced by predominantly negative associations.

**Table 4.**
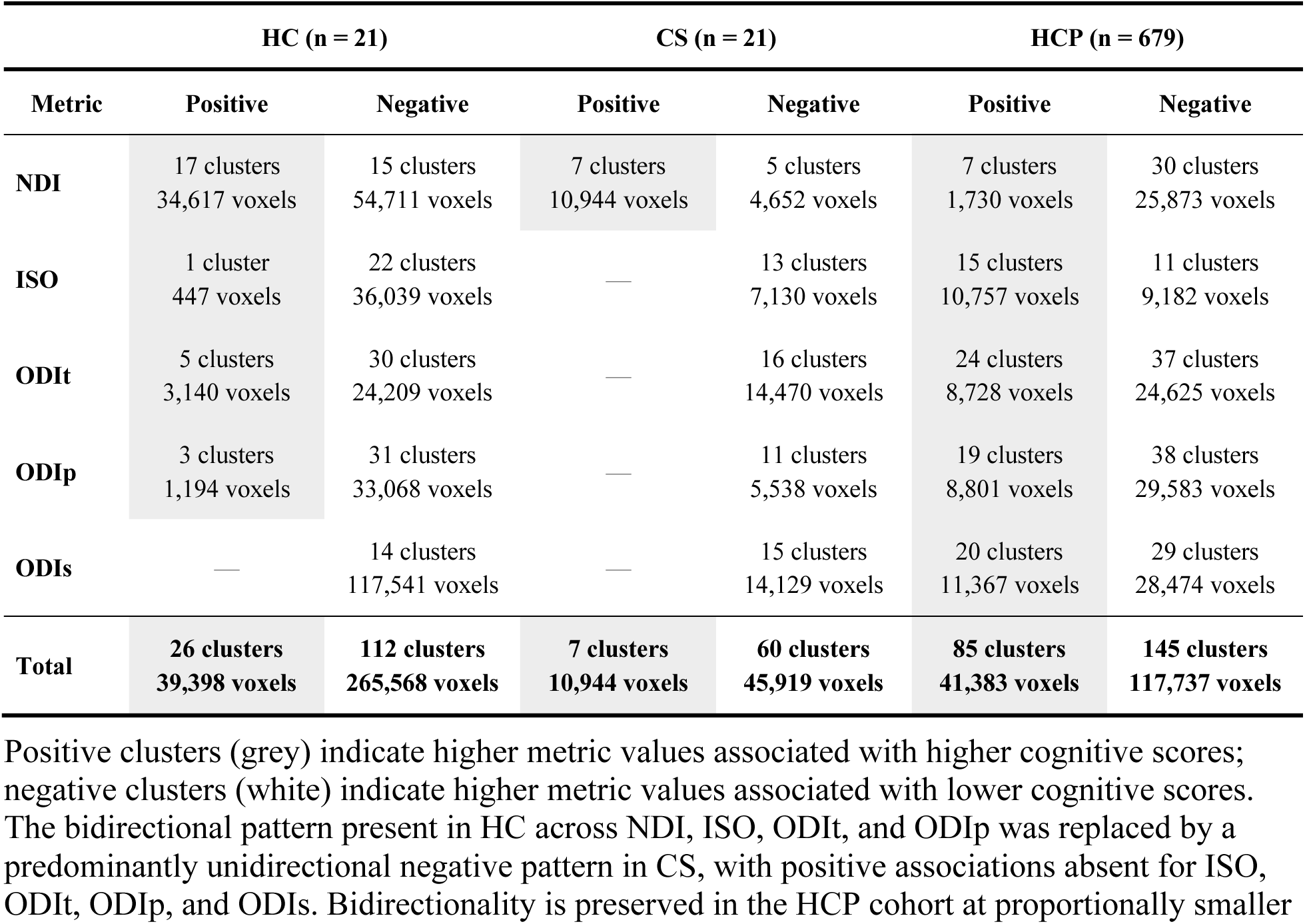
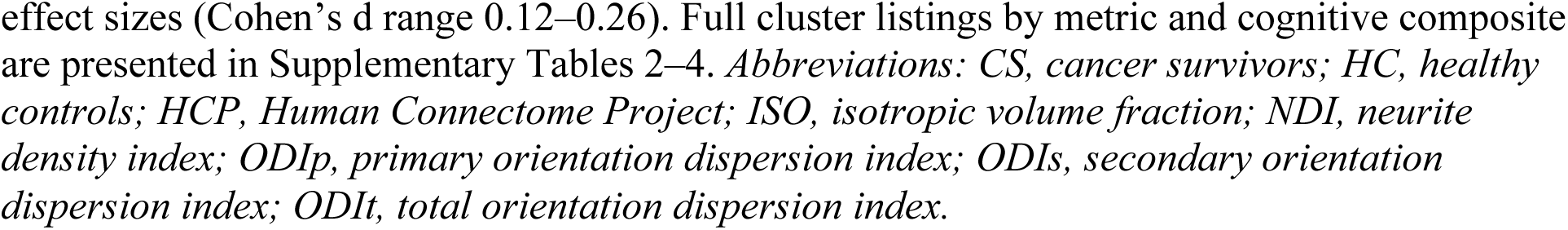
Summary of significant voxel cluster associations between NODDI-Bingham metrics and cognitive performance, by group and direction of association (all three cognitive composites pooled). Counts refer to the number of distinct significant clusters at the p < 0.05 FWE-corrected level; voxel sums are given for each cluster total.

**Table 5.**
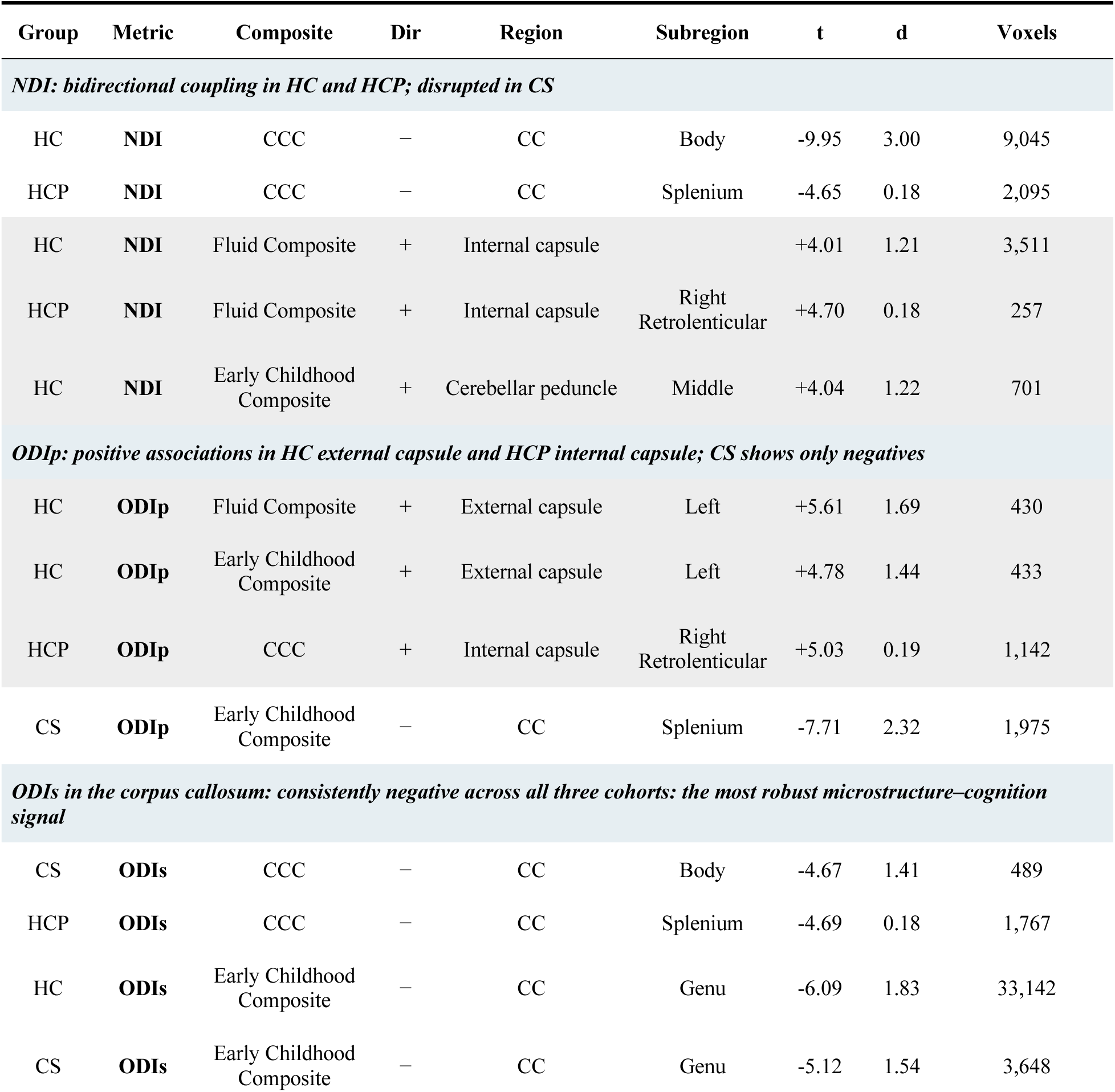
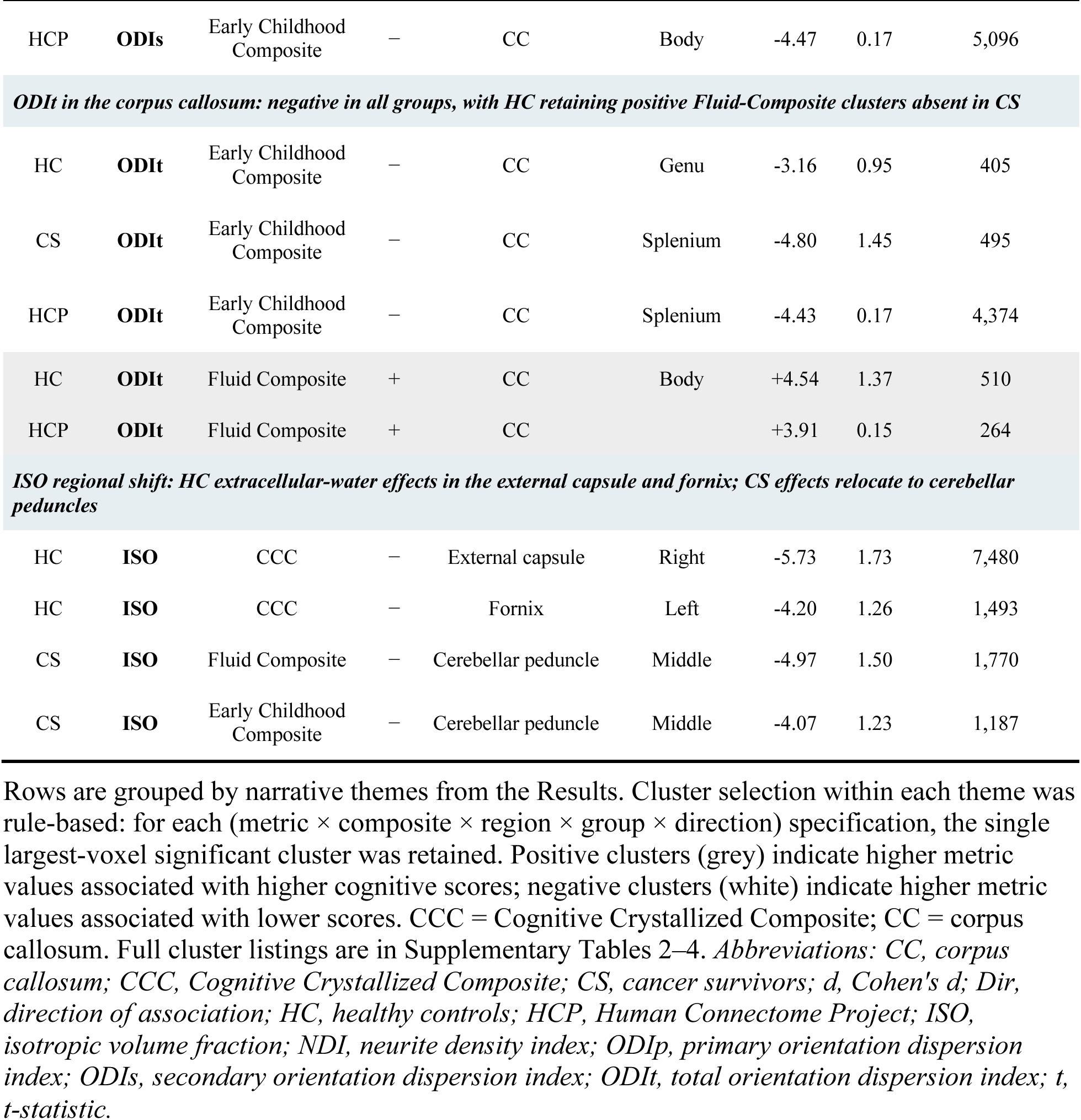
Signature voxel clusters illustrating group-level microstructure–cognition patterns discussed in the Results. For each theme, the largest-voxel cluster matching the specified metric, cognitive composite, anatomical region, group, and direction is reported (p < 0.05 FWE-corrected).

**Figure 3.**
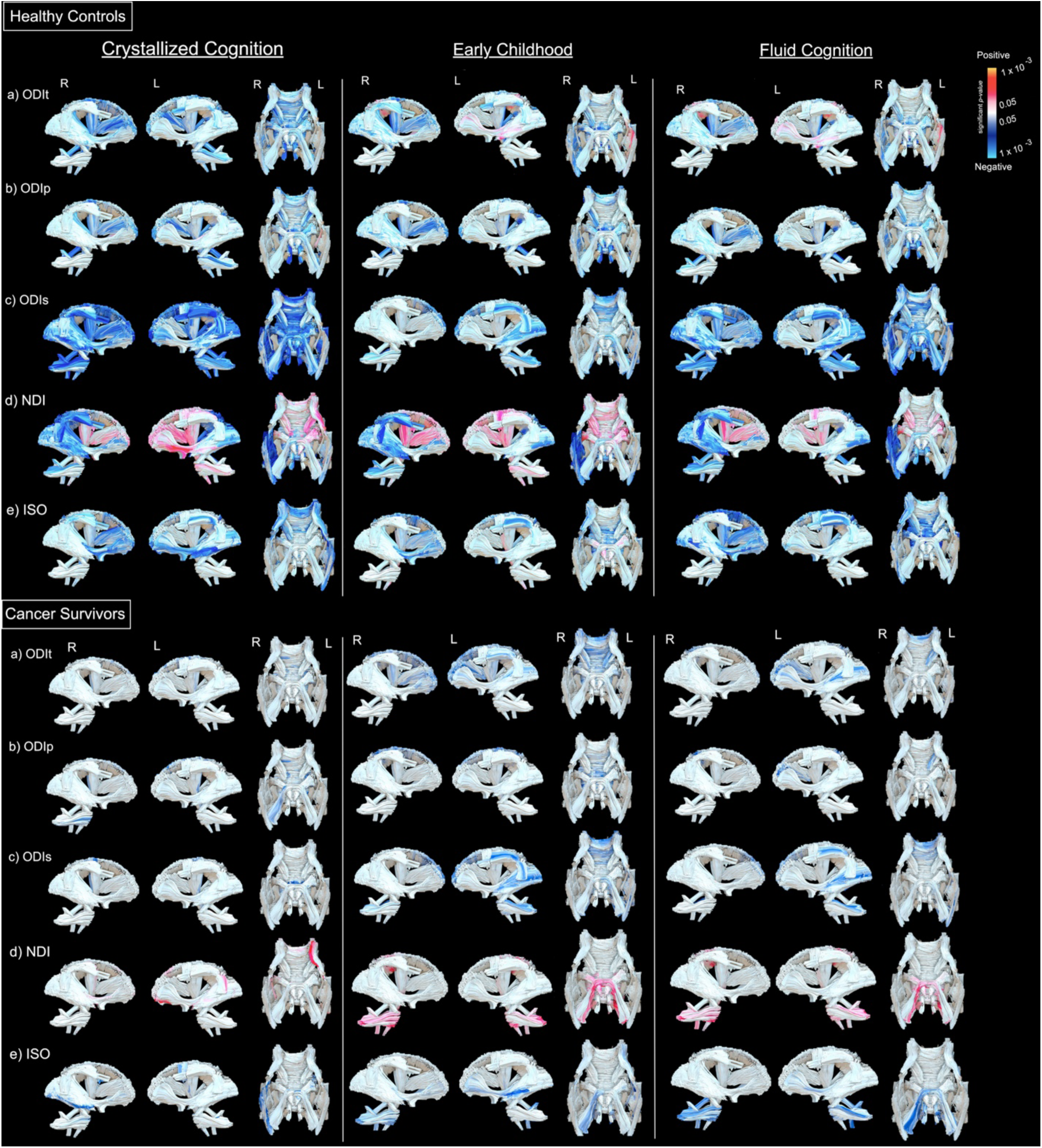
Significant voxelwise regression analyses between NODDI-Bingham white matter metrics and cognitive composite scores in healthy controls (HC; *n* = 21; upper panels) and chemotherapy survivors (CS; *n* = 21; lower panels), covaried for age and sex. Cognitive composites shown are the Cognitive Crystallized Composite (CCC), Fluid Composite, and Early Childhood Composite from the NIH Toolbox. Color indicates significant clusters at *P* < 0.05 FWE-corrected threshold: red indicates positive metric–cognition associations and blue indicates negative associations. HC panels show bidirectional coupling across four of five metrics (NDI, ISO, ODIt, ODIp), with red and blue clusters co-occurring, most strikingly for NDI within the corpus callosum, where positive clusters in the body are accompanied by a strong negative cluster in the splenium (peak *t* = −9.95). ODIs shows exclusively negative associations in HC and the largest spatial extent (117,541 voxels). CS panels show a predominantly unidirectional negative pattern: no positive clusters were detected for ISO, ODIt, ODIp, or ODIs in any cognitive composite. NDI retained reduced bidirectionality in CS (7 positive, 5 negative clusters).

#### Control group

In the HC group, 138 significant clusters spanning 304,966 voxels were observed across all five NODDI-Bingham metrics and all three cognitive composites (Table 4, Fig. 3). Four of the five metrics showed mixed positive and negative relationships with cognition. NDI exhibited this pattern most strikingly within the corpus callosum itself: 17 positive clusters in the CC body, internal capsule, and corona radiata were accompanied by 15 negative clusters (peak *t* = −9.95, *d* = 3.00) concentrated in the CC splenium, thalamic radiation, and cerebellar peduncles, with higher neurite density relating to better cognition in one CC subregion and worse cognition in another. ODIt (5 positive / 30 negative) and ODIp (3 positive / 31 negative) showed analogous mixed patterns, with positive clusters localized to the external capsule and, for ODIt, the CC splenium for the Fluid and Early Childhood Composites. ODIs in the corpus callosum showed the largest spatial extent of any metric (14 clusters, 117,541 voxels, all negative), establishing CC secondary-axis dispersion as the dominant spatial signal in the HC cohort. ISO associations were predominantly negative across the external capsule, fornix, and corona radiata, with a single positive cluster for the Early Childhood Composite in the left cerebral peduncle.

#### Chemotherapy group

In the CS group, only 67 significant clusters spanning 56,863 voxels were identified, roughly half the cluster count and one-fifth the spatial extent of HC (Table 4, Fig. 3). The structure of these associations differed sharply from that of HC. For ISO, ODIt, ODIp, and ODIs, no positive clusters were observed in any cognitive composite: associations in these four metrics were exclusively negative (13, 16, 11, and 15 negative clusters, respectively). Only NDI retained both directions, with 7 positive clusters (cerebellar peduncle and CC, associated with Fluid and Early Childhood Composites) and 5 negative clusters (external capsule, longitudinal fasciculus, corona radiata, and CC, associated with the CCC), but the richer within-CC mixed pattern seen in HC was not present. ISO associations also shifted anatomically, relocating from the external capsule and fornix seen in HC to the cerebellar peduncle, internal capsule, and sagittal stratum. Together, these findings indicate that chemotherapy does not simply attenuate the magnitude of microstructure–cognition relationships but alters the regions and direction of associations.

### HCP Confirmatory Analyses

To assess the generalizability of the HC microstructure–cognition patterns to a large normative sample, voxel-wise regressions were performed in the age-matched HCP subsample (*n* = 679, ages 22–30; Table 5, Fig. 4). Two hundred thirty significant clusters were identified, with effect sizes an order of magnitude smaller than in the primary HC cohort (Cohen’s *d* range 0.12–0.26), consistent with the much larger sample. The mixed relationships observed in HC replicated across every NODDI-Bingham metric: positive and negative clusters were present for NDI (7+/30−), ISO (15+/11−), ODIt (24+/37−), ODIp (19+/38−), and ODIs (20+/29−). Anatomical convergence with HC was strongest for NDI, with positive associations in the internal capsule, corona radiata, and CC across all three cognitive composites in both cohorts, and for ODIs in the corpus callosum, where negative associations were consistently observed across HC, CS, and HCP. ISO showed substantial positive associations with the CCC in the HCP cohort (internal capsule, cerebellar peduncle, CC body) that were largely absent in the primary HC group, consistent with power-related detection limits in the small sample. That these positive associations emerged robustly at *n* = 679 but were absent in the similarly sized CS cohort (*n* = 21) supports the interpretation that the strong negative association pattern in CS reflects genuine chemotherapy-related disruption rather than a small-sample artifact.

**Figure 4.**
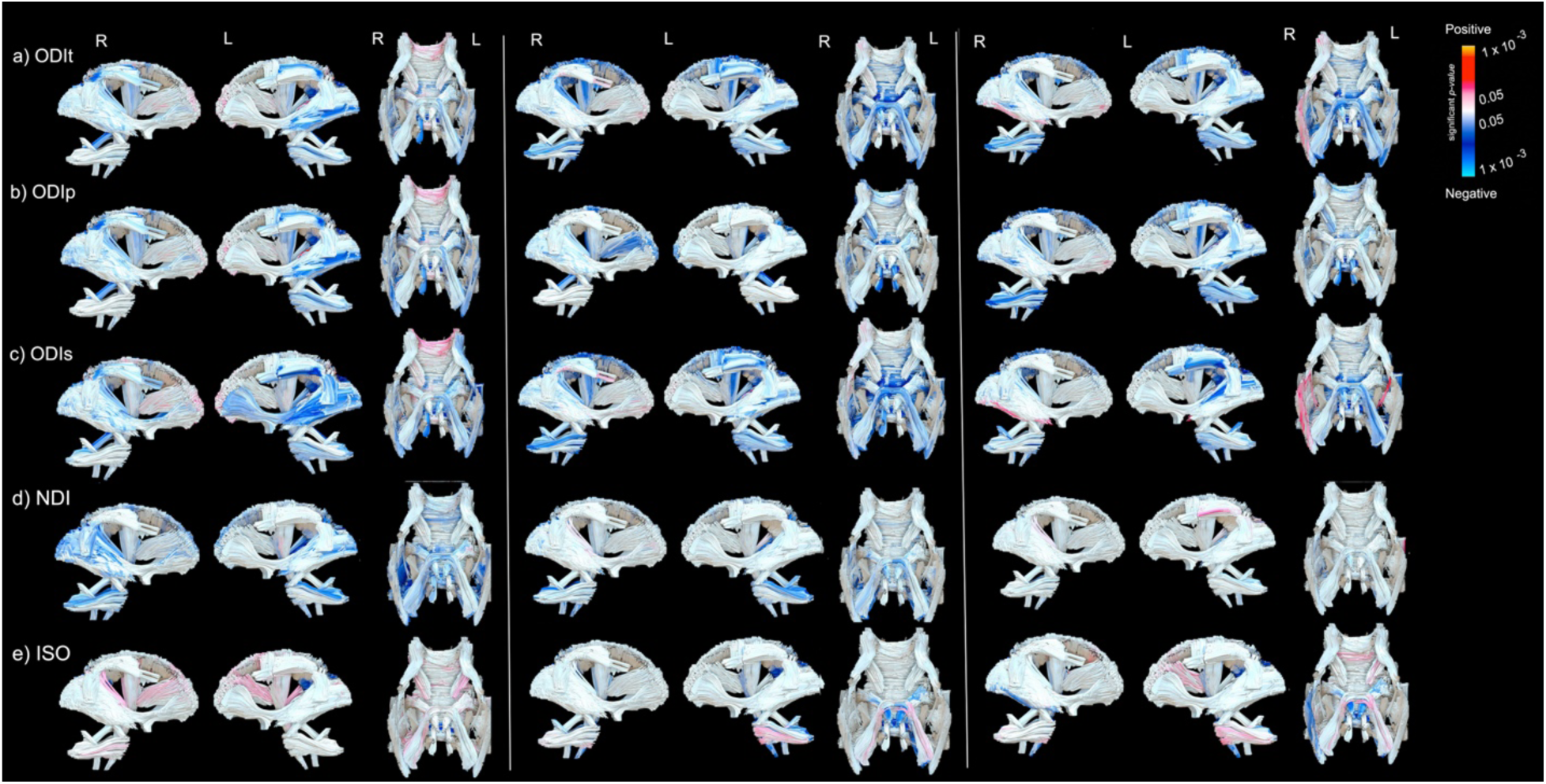
Significant voxelwise regression analyses between NODDI-Bingham white matter metrics and cognitive composite scores in the age-matched HCP subsample (*n* = 679, ages 22–30), covaried for age and sex. Color indicates significant clusters at *P* < 0.05 FWE-corrected threshold: red indicates positive associations and blue indicates negative associations. All five metrics show bidirectional coupling with cognition in this normative cohort, at effect sizes (Cohen’s *d* range 0.12–0.26) consistent with the much larger sample size. Negative associations predominate for orientation dispersion indices (ODIt, ODIp, ODIs) in the corpus callosum splenium and body across all three composites, converging with the HC findings in the primary cohort. Positive associations appear for NDI in the internal capsule and corona radiata, and for ISO and ODIt in multiple tracts. The preserved bidirectional architecture in HCP, in contrast to the unidirectional CS pattern, supports the interpretation that the loss of positive microstructure–cognition coupling in survivors reflects chemotherapy-related disruption rather than a small-sample artifact.

## Discussion

### Group-level microstructural differences: biophysical specificity beyond anisotropy

The group-level microstructural differences observed here localized primarily to cerebellar and brainstem pathways. Survivors showed elevated ODIt and ODIp in the middle cerebellar peduncle and reduced NDI in the corona radiata, cerebral peduncle, and corpus callosum body (Table 3, Fig. 2). These anatomical findings are consistent with prior DTI studies that reported reduced fractional anisotropy in cerebellar and long-range pathways of pediatric cancer survivors^10,12,13^, but NODDI-Bingham’s biophysical decomposition specifies mechanisms that anisotropy alone cannot resolve. The cerebellar signal reflects increased fanning of primary fiber populations (captured by ODIp) rather than secondary-axis dispersion or axonal loss; the corona radiata and callosal signal reflects reduced neurite density rather than fiber disorganization (Fig. 1). These tracts are critical for the sensorimotor integration and cognitive coordination supported by cerebro-cerebellar circuitry through the cerebellar peduncles^28^ and for the interhemispheric communication supported by the corpus callosum^29^, functions most commonly affected in CRCI.

Comparison with our prior IVIM-FWI analysis of this partially overlapping cohort^14^ illustrates the biophysical specificity that NODDI-Bingham adds to anisotropy-based measures. In the middle cerebellar peduncle, IVIM-FWI detected reduced free-water-corrected tissue anisotropy (FAt) in survivors (Cohen’s *d* = 1.43) without resolving the underlying mechanism; NODDI-Bingham attributes this signal specifically to increased primary-axis fiber dispersion (ODIp, *d* = 1.45) and total fiber dispersion (ODIt, peak *d* = 1.42) rather than axonal loss, identifying fanning rather than reduced density as the microstructural signature of cerebellar injury in this cohort. The corpus callosum body illustrates an even clearer case of biophysical resolution. IVIM-FWI reported paradoxically *higher* FAt in survivors than controls in this region (*d* = 1.34), a finding difficult to interpret through anisotropy alone: chemotherapy-exposed white matter should not appear more coherent than healthy white matter. NODDI-Bingham resolves the paradox by showing that neurite density is simultaneously *reduced* in the same region (*d* = 0.93–0.94). The combination, fewer axons packed more coherently, is consistent with selective axonal loss in which surviving fibers retain or increase their geometric alignment. FA-based metrics conflate these two opposing microstructural processes into a single ambiguous signal; NODDI-Bingham’s separate estimation of density and dispersion makes the underlying mechanism interpretable.

Taken together, the group-level findings identify two biologically distinct signatures of chemotherapy-exposed white matter in this cohort: a cerebellar fanning signature captured by ODIp and ODIt, and a cortical-projection axonal loss signature captured by NDI. These are specific, interpretable mechanisms rather than generic reductions in integrity that a DTI-only analysis may have mischaracterized. The question that follows is whether these microstructural differences map onto cognitive deficits, and if so, whether the mapping resembles the relationships observed in healthy white matter.

Whereas group differences localized to cerebellar and brainstem pathways, the microstructure–cognition relationships in HC recruited a broader distributed network including internal capsule, external capsule, corpus callosum body and splenium, and thalamic radiation, regions where CS and HC microstructural values did not differ at the group level. The white matter regions most affected by chemotherapy, in other words, are not identical to the regions through which white matter variation supports cognition in healthy brains. This dissociation indicates that the cognitive consequences of chemotherapy cannot be read off from focal microstructural damage alone; they depend on how well the broader distributed network that supports cognition has survived treatment.

### Mixed positive and negative microstructure–cognition relationships in healthy white matter and HCP replication

In healthy controls, four of five NODDI-Bingham metrics showed a mix of positive and negative microstructure–cognition relationships, with 138 significant clusters spanning approximately 305,000 voxels. The pattern was most striking for NDI, which exhibited 17 positive clusters concentrated in the CC body, internal capsule, and corona radiata alongside 15 negative clusters (peak *t* = −9.95, *d* = 3.00) in the CC splenium, thalamic radiation, and cerebellar peduncles. Higher neurite density therefore related to *better* cognition in one callosal subregion and *worse* cognition in another. ODIt and ODIp showed analogous mixed patterns, with positive clusters localized to the external capsule and CC splenium for the Fluid and Early Childhood Composites. This structure is biologically informative: healthy white matter does not support cognition through a uniform “more is better” rule. The cognitive relevance of a given microstructural value depends on the tract in which it is measured, implying that adaptive fiber arrangements, sheath organization, and axonal packing contribute to cognition through region-specific, context-dependent mechanisms.

Confirmatory analyses in an age-matched HCP subsample (*n* = 679, ages 22–30) supported the interpretation that the HC pattern reflects a genuine feature of healthy white matter rather than an artifact of the primary cohort’s modest sample size. The HCP data reproduced mixed positive and negative microstructure–cognition relationships across all five NODDI-Bingham metrics at proportionally smaller effect sizes (Cohen’s *d* range 0.12–0.26), replicating the positive NDI–cognition associations in the internal capsule, corona radiata, and corpus callosum across all three cognitive composites (Table 5, Fig. 4). One notable extension emerged for ISO, where the larger HCP sample detected positive ISO–cognition associations in the internal capsule, cerebellar peduncle, and CC body that were largely undetectable in the smaller HC group, suggesting that healthy ISO–cognition relationships have their own mixed structure that was masked by power in the primary cohort. This pattern of replication with appropriate effect-size attenuation establishes the HC findings as a normative reference against which survivor relationships can be compared.

### Loss of positive relationships in cancer survivors: a candidate mechanism for CRCI

Against this HC/HCP normative backdrop, survivors showed a qualitatively different pattern. Across 67 clusters spanning approximately 57,000 voxels, roughly half the cluster count and one-fifth the spatial extent of HC, four of five metrics lost their positive clusters entirely. ISO, ODIt, ODIp, and ODIs showed exclusively negative relationships with cognition in CS. Only NDI retained a reduced form of its mixed pattern (7 positive and 5 negative clusters), but the rich within-CC signature seen in HC, simultaneous body-positive and splenium-negative associations, did not survive. For ODIt and ODIp, the loss was categorical: 0 positive clusters in CS against 5 and 3 positive clusters in HC. The architecture characterizing healthy microstructure–cognition relationships was not merely attenuated; it was replaced by a uniformly negative pattern.

The restructuring is biologically interpretable through the metrics that changed. The reduction in NDI in the corona radiata, cerebral peduncles, and corpus callosum of CS is consistent with the well-documented toxicity of chemotherapeutic agents to oligodendrocytes and their precursors^6,30^, which would be expected to reduce the effective axonal volume fraction in affected tracts. The categorical loss of positive ODIp and ODIt clusters is consistent with disruption of the fiber-arrangement coherence that depends on intact myelin sheath organization^18^: in healthy white matter, modest increases in primary-axis dispersion in particular tracts appear to carry cognitive information, whereas in chemotherapy-exposed white matter, the same microstructural variation no longer does. Whether this represents loss of an active regulatory capacity, residual heterogeneity from tract-specific injury patterns, or some combination cannot be distinguished with the present data. What is clear is that the healthy-white-matter pattern, in which multiple metrics each relate to cognition in both directions depending on region, is specifically absent in survivors.

One metric–region relationship survived the chemotherapy-related restructuring intact: ODIs in the corpus callosum remained negatively associated with cognition in every group examined, with 14 clusters in HC, 15 in CS, and 29 in the HCP subsample. Both the directionality and the concentration in callosal regions were preserved in survivors, with the HC/CS difference in spatial extent (∼8-fold) reflecting the overall reduction in the number of significant relationships rather than a shift in the underlying relationship itself. That ODIs survived while NDI, ODIt, and ODIp did not points to a specific biological distinction: secondary-axis dispersion in callosal tracts may reflect a baseline architectural property of the tract that is comparatively resistant to chemotherapy-related disruption. This resistance identifies callosal ODIs as a candidate cross-population biomarker of cognitive function that remains interpretable across healthy and chemotherapy-exposed white matter.

### Regional shift in ISO–cognition relationships

ISO–cognition associations shifted anatomically between groups rather than disappearing. In HC, they were concentrated in the external capsule, fornix, and thalamic radiation; in CS, they relocated to the cerebellar peduncle, internal capsule, and sagittal stratum. This regional divergence suggests that chemotherapy differentially affects the extracellular environment across white matter tracts, potentially reflecting tract-specific vulnerability to neuroinflammation and edema, processes known to accompany chemotherapy-induced oligodendrocyte injury. HCP data added a note of interest: positive ISO–cognition associations emerged with the larger sample across the internal capsule, cerebellar peduncle, and CC body, a pattern largely undetectable in the small HC group. This suggests that healthy ISO–cognition relationships have their own mixed positive and negative structure that was masked by power in the primary cohort, and that the CS regional relocation may represent both a shift and a loss of the positive component. Furthermore, although ISO showed no group-level difference between survivors and controls, it remained reliably coupled to cognition. This dissociation parallels our IVIM-FWI analysis of the overlapping cohort, in which the model’s free-water fraction was elevated in survivors across multiple white matter tracts and was itself associated with cognitive performance^14^.

## Limitations and Future Directions

Several limitations should be considered. The modest sample size (*n* = 21 per group) may have limited power to detect smaller effects, particularly positive microstructure–cognition associations in HC, as evidenced by the broader mixed positive and negative patterns recovered in the HCP cohort. The cross-sectional design precludes causal inference regarding whether the observed microstructural differences preceded or followed cognitive decline. The heterogeneity of cancer diagnoses and treatment regimens within the CS group may have introduced variability, although this also enhances generalizability. Future longitudinal studies with larger survivor samples should examine whether NODDI-Bingham metrics predict cognitive trajectories over time and whether the disruption of microstructure–cognition relationships is progressive, stable, or recovers after treatment cessation. Interventional studies targeting oligodendrocyte health in survivors could test whether preserving or restoring the mixed positive and negative pattern improves cognitive outcomes. Myelin-targeting agents such as clemastine and other remyelination-promoting compounds are being investigated in populations ranging from multiple sclerosis to traumatic brain injury^31,32^. The metric specificity of NODDI-Bingham provides a more precise target engagement readout than FA-based imaging: agents acting on myelin sheath organization would be expected to modulate ODIp, while interventions promoting axonal preservation would primarily affect NDI. This capacity to dissociate myelin from axonal effects at the imaging level could meaningfully inform the design of neuroprotective trials in pediatric cancer survivors, where rational selection of primary endpoints depends on matching the biomarker to the drug’s mechanism of action.

## Conclusion

This study demonstrates that NODDI-Bingham reveals aspects of chemotherapy-related white matter pathology that conventional DTI cannot: not only reduced neurite density and increased fiber dispersion, but also a restructuring of how white matter microstructure relates to cognition. The preservation of ODIs–cognition associations across healthy controls, cancer survivors, and a large normative cohort identifies corpus callosum secondary-axis dispersion as a cross-population marker of cognitive function. The specific loss of positive NDI, ODIt, and ODIp relationships in survivors points to disrupted regulatory microstructural mechanisms as a candidate mechanism for chemotherapy-related cognitive change. Together, these findings highlight NODDI-Bingham as a sensitive biomarker panel for monitoring white matter health in pediatric cancer survivors and for evaluating neuroprotective interventions targeting the relationships between white matter structure and cognitive function.

## Data Availability Statement

The individual-level diffusion MRI and cognitive data from the pediatric cancer survivor and healthy control cohorts are not publicly available due to the sensitive nature of pediatric patient data and institutional restrictions but are available from the corresponding author upon reasonable request and subject to a data-use agreement. The Human Connectome Project Young Adult data used for normative replication are publicly available through ConnectomeDB (https://db.humanconnectome.org). Analyses used the dmipy toolbox for NODDI-Bingham fitting^23^.

## Acknowledgements

We greatly thank all of our participants and their families for their help with this study and the MRI staff at Albert Einstein College of Medicine Magnetic Resonance Imaging Research Center for their MRI acquisition efforts. Data were provided in part by the Human Connectome Project, WU-Minn Consortium (Principal Investigators: David Van Essen and Kamil Ugurbil; 1U54MH091657) funded by the 16 NIH Institutes and Centers that support the NIH Blueprint for Neuroscience Research; and by the McDonnell Center for Systems Neuroscience at Washington University.

## Author contributions

Tim Q. Duong designed the study. Sonya Henry and Shounak Nandi acquired the data. Shounak Nandi and Faye McKenna analyzed the data. Shounak Nandi, Faye McKenna, and Tim Q. Duong wrote and Siddharth Nayak revised the paper. All authors approved the submitted and final versions of the manuscript.

