## Supplemental Material for "White matter–cognition mapping in pediatric cancer survivors: a two-cohort NODDI-Bingham study"

### 1. Supplementary Methods

*Participants and Eligibility*

Inclusion criteria required that cancer patients be able to undergo MRI without sedation and have received standard chemotherapy protocols. Healthy controls were recruited from the community and matched to cancer participants by age. Because of the limited sample sizes, not all demographic variables were well matched.

Exclusion criteria for all participants included contraindications for MRI; pre-existing neurodegenerative, psychiatric, or neurocognitive disorders; prior or anticipated receipt of brain-directed therapy; and prior hospitalization for critical illness. Brain cancer patients and those requiring sedation for MRI were excluded. Healthy controls were required to provide assent and to be generally healthy.

After application of the inclusion and exclusion criteria and an image-quality review (excluding participants whose diffusion data showed significant motion that could not be re-acquired), 21 cancer survivors (CS) and 21 healthy controls (HC) were retained for analysis (N = 42).

*NIH Toolbox Cognition Battery*

Cognitive performance was assessed using the NIH Toolbox Cognition Battery[1], administered to evaluate multiple cognitive domains. Analyses focused on three composite scores. The Crystallized Cognition Composite (CCC) is derived from the Picture Vocabulary and

Oral Reading Recognition tests and indexes language and verbal reasoning. The Fluid Cognition Composite is based on the Flanker Inhibitory Control and Attention, Dimensional Change Card Sort, Picture Sequence Memory, List Sorting, and Pattern Comparison tests, together assessing problem-solving, adaptability, processing speed, and memory encoding. The Early Childhood Composite summarizes performance on the early-childhood-validated subset of these tasks (Picture Vocabulary, Flanker Inhibitory Control and Attention, Dimensional Change Card Sort, and Picture Sequence Memory).

*Magnetic Resonance Imaging Acquisition*

MRI data were acquired on a 3T Philips Ingenia Elition scanner (Achieva, Philips, Amsterdam, Netherlands) using a 32-element head-neck coil. Three-dimensional T1-weighted (T1w) images were acquired with a magnetization-prepared rapid gradient-echo (MPRAGE) sequence (repetition time [TR] 2.3 s, echo time [TE] 3.24 ms, field of view 224 × 224 × 224 mm, matrix 256 × 256 × 256, 0.87 × 0.87 × 0.87 mm isotropic voxels). Diffusion MRI data were acquired with a multi-band diffusion-weighted spin-echo echo-planar imaging sequence (anterior-to-posterior phase encoding, 2 mm isotropic voxels, TR 1250 ms, TE 93 ms, multi-band factor 2) with b-values = 0 (6 volumes), 300 (6), 800 (27), and 2000 (30) s/mm$^2$. A three-dimensional b0 field map was acquired to correct for distortions (2 mm voxels, TR 2600 ms, TE 94 ms, SENSE multi-band factor 2).

*Diffusion MRI Preprocessing*

Diffusion images were first denoised using MRtrix3 dwidenoise[2] and corrected for Gibbs-ringing artifacts[3]. Images were then corrected for head motion, eddy-current distortions, and susceptibility-induced B0 field inhomogeneities using FSL topup and eddy[4].

*NODDI-Bingham Model*

The multi-compartment NODDI-Bingham model[5] was fit to the preprocessed diffusion data using Diffusion Microstructure Imaging in Python (DMIPY)[6]. Whereas the standard NODDI model uses a Watson distribution that assumes symmetric dispersion about a single axis, the Bingham extension models anisotropic dispersion and so accounts for the fanning and bending of fiber populations. The model yields five microstructural metrics: the neurite density index (NDI), the isotropic free-water volume fraction (ISO), and three orientation dispersion indices. The primary (ODIp) and secondary (ODIs) indices capture dispersion along the two principal axes of the dispersion plane, where ODIp reflects fanning within the plane of greatest dispersion and ODIs reflects dispersion perpendicular to that plane, while the total dispersion index (ODIt) summarizes overall dispersion. The Bingham distribution is governed by a concentration parameter and an anisotropy parameter, the latter equal to zero in the symmetric Watson/NODDI case and increasing with more asymmetric fanning. Full parameter definitions and the corresponding equations are provided in the main-text Methods.

*Whole-Sample Associations Between NODDI-Bingham Metrics and Cognitive Performance*

Voxel-wise regression analyses in the whole sample ($N = 42$), controlling for age, sex and group membership, revealed widespread associations between NODDI-Bingham metrics and cognitive test scores (Supplementary Table 1).

**2. Supplementary Results**

*Neurite density index.* The NDI exhibited bidirectional associations with cognition. Higher NDI in the right posterior thalamic radiation and CC splenium was associated with lower CCC scores (peak $t = -3.61$, $d = 1.09$). Conversely, higher NDI was positively associated with the Fluid Composite across large clusters in the CC body ($t = 4.20$, $d = 1.26$; 3,871 voxels), right posterior corona radiata, left cerebral peduncle, left SLF, and middle cerebellar peduncle. Similar positive associations were observed with the Early Childhood Composite in the right posterior corona radiata, CC body, left cerebral peduncle, and middle cerebellar peduncle.

*Isotropic volume fraction.* Higher ISO was associated with lower scores across all three cognitive composites. Associations with the CCC were observed in the left superior corona radiata ($t = -3.81$, $d = 1.15$; 1,211 voxels). The Fluid Composite showed extensive negative associations with ISO in the right external capsule ($t = -4.61$, $d = 1.39$; 4,913 voxels), middle cerebellar peduncle, left fornix, left posterior thalamic radiation, left SLF, and left retrolenticular internal capsule. The Early Childhood Composite was similarly associated with higher ISO in the right external capsule, left fornix, middle cerebellar peduncle, and left sagittal stratum.

*Total orientation dispersion index.* Higher ODIt was consistently associated with lower cognitive scores across all three composites. The CCC was negatively associated with ODIt in the right superior corona radiata, CC (body and genu), middle cerebellar peduncle, right cerebral peduncle, and right retrolenticular internal capsule (peak $t = -5.03$, $d = 1.51$). The Fluid Composite demonstrated negative associations across ten clusters spanning the CC (splenium, genu, and body), right posterior thalamic radiation, left posterior internal capsule, right posterior corona radiata, right corticospinal tract, and right external capsule. The Early Childhood Composite showed the most widespread ODIt associations, with a large cluster in the CC splenium (10,947 voxels; $t = -6.13$, $d = 1.85$), along with involvement of the posterior thalamic radiation, left posterior internal capsule, and right cerebral peduncle.

*Primary orientation dispersion index.* The ODIp showed a divergent pattern across cognitive domains. Higher ODIp was negatively associated with the CCC in the CC (body and genu), right superior cerebellar peduncle, right superior corona radiata, and right cerebral peduncle (peak $t = -5.28$, $d = 1.59$). Strong negative associations with the Fluid Composite were demonstrated in the CC splenium ($t = -4.61$, $d = 1.39$; 1,765 voxels), left superior and right anterior coronae radiatae, CC body, right inferior cerebellar peduncle, CC genu, and right posterior thalamic radiation. A single positive cluster was found in the left external capsule ($t = 4.6$, $d = 1.39$, 450 voxels). Interestingly, this region showed a positive association with the Early Childhood Composite ($t = 4.79$, $d = 1.44$, 386 voxels). Otherwise, a similar negative pattern was observed with the Early Childhood Composite score: a large cluster in the CC splenium encompassed 2,859 voxels ($t = -6.38$, $d = 1.92$), accompanied by associations in the CC (genu and body), right cerebral peduncle, right anterior corona radiata, right anterior internal capsule, right SLF, and right posterior thalamic radiation.

*Secondary orientation dispersion index.* Higher ODIs was negatively associated with all three cognitive composites across widespread white matter regions. The CCC was negatively associated with ODIs in the CC body (two clusters totaling 7,937 voxels; peak $t = -5.22$, $d = 1.57$), middle cerebellar peduncle, left retrolenticular internal capsule, right cingulum, left SLF, and right posterior corona radiata. The Fluid Composite showed the most extensive ODIs associations, with large clusters in the CC splenium (8,650 voxels; $t = -4.59$, $d = 1.38$) and CC genu (6,534 voxels), as well as in the middle cerebellar peduncle, right cerebral peduncle, and right SLF. The Early Childhood Composite was negatively associated with ODIs across the largest spatial extent of any metric, including the left SLF (22,398 voxels; $t = -5.31$, $d = 1.60$), CC splenium (15,210 voxels; $t = -6.43$, $d = 1.94$), and middle cerebellar peduncle.

### 3. Supplementary Tables

**Supplementary Table 1** Significant voxel cluster associations between NODDI-Bingham metrics and cognitive scores in the whole sample at the $P < 0.05$ FWE-corrected level.

| NODDI metric | Cognitive test | Voxels | Peak t-stat | P-Value | Peak Cohen's d | Coordinates [X Y Z] | Region | Subregion |
|---|---|---|---|---|---|---|---|---|
| **NDI** | ***CCC*** | 971 | -3.25 | <0.001 | 0.98 | [56 74 77] | Thalamic radiation | Right Posterior |
| | | 780 | -3.61 | <0.001 | 1.09 | [90 87 85] | CC | Splenium |
| | | 657 | -3.54 | <0.001 | 1.07 | [114 73 83] | | Splenium |
| | ***Fluid Composite*** | 3871 | 4.2 | <0.001 | 1.26 | [97 118 101] | CC | Body |
| | | 3262 | 4.09 | <0.001 | 1.23 | [66 101 102] | Corona radiata | Right Posterior |
| | | 1235 | 4.4 | <0.001 | 1.33 | [102 100 54] | Cerebral peduncle | Left |
| | | 902 | 3.37 | <0.001 | 1.01 | [125 88 100] | Longitudinal fasciculus | Left Superior |
| | | 578 | 3.61 | 0.004 | 1.09 | [75 96 44] | Cerebellar peduncle | Middle |
| | | 506 | 3.64 | 0.013 | 1.1 | [105 80 87] | CC | Splenium |
| | | 670 | -3.51 | 0.001 | 1.06 | [56 81 79] | Thalamic radiation | Right Posterior |
| | ***Early Childhood Composite*** | 1443 | 3.98 | <0.001 | 1.2 | [66 100 102] | Corona radiata | Right Posterior |
| | | 1327 | 4.32 | <0.001 | 1.3 | [97 118 101] | CC | Body |
| | | 1120 | 4.2 | <0.001 | 1.26 | [101 112 58] | Cerebral peduncle | Left |
| | | 913 | 4.1 | <0.001 | 1.23 | [76 97 45] | Corona radiata | Right Superior |
| | | 2136 | -4.6 | <0.001 | 1.39 | [56 81 79] | Thalamic radiation | Right Posterior |
| | | 948 | -3.53 | <0.001 | 1.06 | [45 115 94] | Longitudinal fasciculus | Right Superior |
| **ISO** | ***CCC*** | 1211 | -3.81 | <0.001 | 1.15 | [117 133 95] | Corona radiata | Left Superior |
| | | 522 | -4.36 | <0.001 | 1.31 | [125 129 71] | Internal capsule | |
| | ***Fluid Composite*** | 4913 | -4.61 | <0.001 | 1.39 | [58 127 65] | External capsule | Right |
| | | 2101 | -3.87 | <0.001 | 1.17 | [117 63 35] | Cerebellar peduncle | |
| | | 1929 | -4.27 | <0.001 | 1.29 | [66 83 39] | Cerebellar peduncle | Middle |
| | | 791 | -4.87 | <0.001 | 1.47 | [122 115 58] | Fornix | Left |
| | | 522 | -3.39 | 0.004 | 1.02 | [128 81 73] | Thalamic radiation | Left Posterior |
| | | 496 | -3.80 | 0.006 | 1.14 | [125 114 98] | Longitudinal fasciculus | Left Superior |
| | | 467 | -3.53 | 0.010 | 1.06 | [118 103 83] | Internal capsule | Left Retrolenticular |

| | | | | | | | | |
|---|---|---|---|---|---|---|---|---|
| | ***Early Childhood Composite*** | 3155 | -4.38 | <0.001 | 1.32 | [58 127 65] | External capsule | Right |
| | | 1414 | -5.03 | <0.001 | 1.51 | [122 116 57] | Fornix | Left |
| | | 1342 | -4.58 | <0.001 | 1.38 | [65 85 40] | Cerebellar peduncle | Middle |
| | | 528 | -3.37 | 0.001 | 1.01 | [112 75 37] | | Middle |
| | | 491 | -3.25 | 0.002 | 0.98 | [130 92 66] | Sagittal stratum | Left |
| | | 437 | -3.66 | 0.005 | 1.10 | [110 67 41] | Cerebellar peduncle | Middle |
| **ODIt** | ***CCC*** | 973 | -4.73 | <0.001 | 1.42 | [71 108 108] | Corona radiata | Right Superior |
| | | 851 | -4.41 | <0.001 | 1.33 | [105 123 109] | CC | Body |
| | | 684 | -4.33 | <0.001 | 1.30 | [70 75 43] | Cerebellar peduncle | Middle |
| | | 606 | -4.48 | <0.001 | 1.35 | [99 150 87] | CC | Genu |
| | | 344 | -5.03 | 0.003 | 1.51 | [83 110 57] | Cerebral peduncle | Right |
| | | 275 | -4.04 | 0.020 | 1.22 | [56 88 77] | Internal capsule | Right Retrolenticular |
| | ***Fluid Composite*** | 967 | -5.37 | <0.001 | 1.62 | [104 88 95] | CC | Splenium |
| | | 627 | -3.63 | <0.001 | 1.09 | [87 150 79] | | Genu |
| | | 484 | -4.85 | <0.001 | 1.46 | [55 74 72] | Thalamic radiation | Right Posterior |
| | | 477 | -5.11 | <0.001 | 1.54 | [106 103 56] | Cerebral peduncle | |
| | | 464 | -3.85 | <0.001 | 1.16 | [105 122 83] | Internal capsule | Left Posterior |
| | | 461 | -3.54 | <0.001 | 1.07 | [67 100 95] | Corona radiata | Right Posterior |
| | | 401 | -4.28 | 0.001 | 1.29 | [82 101 49] | Corticospinal tract | Right |
| | | 360 | -3.74 | 0.002 | 1.13 | [72 108 107] | CC | Body |
| | | 320 | -4.26 | 0.005 | 1.28 | [57 121 68] | External capsule | Right |
| | | 295 | -4.39 | 0.010 | 1.32 | [74 92 103] | CC | Splenium |
| | ***Early Childhood Composite*** | 10947 | -6.13 | <0.001 | 1.85 | [74 92 103] | CC | Splenium |
| | | 1700 | -5.28 | <0.001 | 1.59 | [55 73 73] | Thalamic radiation | Right Posterior |
| | | 541 | -4.95 | <0.001 | 1.49 | [106 103 56] | Cerebral peduncle | |
| | | 514 | -4.17 | <0.001 | 1.26 | [105 121 82] | Internal capsule | Left Posterior |
| | | 448 | -4.53 | 0.002 | 1.36 | [72 104 65] | Cerebral peduncle | Right |
| | | 441 | -4.27 | 0.002 | 1.29 | [94 91 79] | CC | |
| | | 393 | -4.55 | 0.007 | 1.37 | [126 80 83] | Thalamic radiation | Left Posterior |
| **ODIp** | ***CCC*** | 966 | -4.37 | <0.001 | 1.32 | [72 111 107] | CC | Body |
| | | 795 | -5.28 | <0.001 | 1.59 | [106 109 104] | | Body |
| | | 646 | -4.23 | <0.001 | 1.27 | [83 77 44] | Cerebellar peduncle | Right Superior |
| | | 513 | -4.75 | <0.001 | 1.43 | [89 148 80] | CC | Genu |
| | | 410 | -4.55 | <0.001 | 1.37 | [74 124 112] | Corona radiata | Right Superior |
| | | 292 | -4.7 | 0.01 | 1.42 | [83 111 57] | Cerebral peduncle | Right |
| | ***Fluid Composite*** | 450 | 4.6 | <0.001 | 1.39 | [122 130 74] | External capsule | Left |

| | | | | | | | | |
|---|---|---|---|---|---|---|---|---|
| | | 1765 | -4.61 | <0.001 | 1.39 | [99 89 79] | CC | Splenium |
| | | 544 | -4.91 | <0.001 | 1.48 | [116 103 97] | Corona radiata | Left Superior |
| | | 400 | -3.96 | 0.001 | 1.19 | [72 109 108] | CC | Body |
| | | 343 | -4.29 | 0.004 | 1.29 | [79 82 43] | Cerebellar peduncle | Right Inferior |
| | | 332 | -4.3 | 0.006 | 1.3 | [84 148 68] | CC | Genu |
| | | 286 | -4.24 | 0.02 | 1.28 | [55 74 72] | Thalamic radiation | Right Posterior |
| | | 285 | -3.66 | 0.021 | 1.1 | [71 156 67] | Corona radiata | Right Anterior |
| | ***Early Childhood Composite*** | 386 | 4.79 | 0.004 | 1.44 | [124 126 69] | External capsule | Left |
| | | 2859 | -6.38 | <0.001 | 1.92 | [75 92 102] | CC | Splenium |
| | | 1724 | -4.7 | <0.001 | 1.42 | [103 89 95] | | Splenium |
| | | 1433 | -4.47 | <0.001 | 1.35 | [84 148 68] | | Genu |
| | | 1376 | -5.21 | <0.001 | 1.57 | [108 108 105] | | Body |
| | | 428 | -4.58 | 0.002 | 1.38 | [72 104 65] | Cerebral peduncle | Right |
| | | 409 | -3.59 | 0.002 | 1.08 | [68 158 82] | Corona radiata | Right Anterior |
| | | 337 | -4.31 | 0.014 | 1.3 | [75 132 77] | Internal capsule | Right Anterior |
| | | 320 | -4.69 | 0.021 | 1.41 | [52 95 107] | Longitudinal fasciculus | Right Superior |
| | | 319 | -4.71 | 0.022 | 1.42 | [55 74 73] | Thalamic radiation | Right Posterior |
| **ODIs** | ***CCC*** | 4675 | -5.22 | <0.001 | 1.57 | [73 107 107] | CC | Body |
| | | 3262 | -4.12 | <0.001 | 1.24 | [101 125 103] | | Body |
| | | 1655 | -4.02 | <0.001 | 1.21 | [69 92 41] | Cerebellar peduncle | Middle |
| | | 796 | -3.79 | <0.001 | 1.14 | [126 88 76] | Internal capsule | Left Retrolenticular |
| | | 746 | -3.42 | <0.001 | 1.03 | [81 158 86] | Cingulum | Right |
| | | 614 | -3.28 | 0.001 | 0.99 | [130 114 101] | Longitudinal fasciculus | Left Superior |
| | | 470 | -3.96 | 0.013 | 1.19 | [57 77 92] | Corona radiata | Right Posterior |
| | ***Fluid Composite*** | 8650 | -4.59 | <0.001 | 1.38 | [105 88 103] | CC | Splenium |
| | | 7723 | -5.23 | <0.001 | 1.58 | [73 89 103] | | Splenium |
| | | 6534 | -4.34 | <0.001 | 1.31 | [93 155 75] | | Genu |
| | | 2736 | -4.42 | <0.001 | 1.33 | [105 101 55] | Cerebral peduncle | |
| | | 1525 | -3.97 | <0.001 | 1.20 | [67 84 36] | Cerebellar peduncle | Middle |
| | | 1506 | -4.25 | <0.001 | 1.28 | [79 107 57] | Cerebral peduncle | Right |
| | | 855 | -3.81 | <0.001 | 1.15 | [54 110 103] | Longitudinal fasciculus | Right Superior |
| | | 569 | -3.80 | 0.015 | 1.14 | [131 106 63] | External capsule | |
| | ***Early Childhood Composite*** | 22398 | -5.31 | <0.001 | 1.60 | [128 113 102] | Longitudinal fasciculus | Left Superior |
| | | 15210 | -6.43 | <0.001 | 1.94 | [73 89 103] | CC | Splenium |
| | | 3829 | -4.51 | <0.001 | 1.36 | [110 71 39] | Cerebellar peduncle | Middle |
| | | 2190 | -4.46 | <0.001 | 1.34 | [72 66 38] | | Middle |

Grey cells indicate a positive t-statistic and white cells indicate a negative t-statistic. Significance was determined using a permutation cluster-wise $P < 0.050$ familywise error (FWE)-corrected threshold. Positive t-statistic values indicate a significantly positive regression between the diffusion signal and cognitive test, while negative t-statistic values indicate a significantly negative regression between the diffusion signal and cognitive test in the whole group (CS n = 21, HC n = 21). *Abbreviations: [X Y Z], coordinates of the voxels on the MNI 1 mm brain atlas; CC, corpus callosum; CCC, Cognitive Crystallized Composite; CS, cancer survivors; FWE, familywise error; HC, healthy controls; HCP, Human Connectome Project; ISO, isotropic volume fraction; NDI, neurite density index; ODIp, primary orientation dispersion index; ODIs, secondary orientation dispersion index; ODIt, total orientation dispersion index; t, t-statistic; WM, white matter.*

**Supplementary Table 2** Significant voxel cluster associations between NODDI-Bingham metrics and cognitive scores in the HC group at the $P < 0.05$ FWE-corrected level.

| NODDI metric | Cognitive test | Voxels | Peak t-stat | P-Value | Peak Cohen's d | Coordinates [X Y Z] | Region | Subregion |
|---|---|---|---|---|---|---|---|---|
| **NDI** | ***CCC*** | 9027 | 4.82 | <0.001 | 1.45 | [109 131 82] | Internal capsule | Left Anterior |
| | | 1994 | 4.58 | <0.001 | 1.38 | [60 114 86] | External capsule | Right |
| | | 785 | 5.56 | <0.001 | 1.67 | [126 120 59] | Internal capsule | |
| | | 602 | 5.19 | 0.001 | 1.56 | [81 119 72] | CC | |
| | | 595 | 5.06 | 0.001 | 1.52 | [74 157 78] | CC | Genu |
| | | 545 | 5.14 | 0.003 | 1.55 | [101 102 38] | Cerebellar peduncle | Middle |
| | | 13206 | -5.61 | <0.001 | 1.69 | [68 95 106] | Corona radiata | Right Posterior |
| | | 9045 | -9.95 | <0.001 | 3.00 | [79 98 102] | CC | Body |
| | | 1339 | -4.46 | <0.001 | 1.34 | [73 80 42] | Cerebellar peduncle | Middle |
| | | 777 | -5.69 | <0.001 | 1.71 | [70 154 73] | CC | |
| | | 433 | -4.35 | 0.020 | 1.31 | [84 89 43] | Medial lemniscus | Right |
| | ***Fluid Composite*** | 3511 | 4.01 | <0.001 | 1.21 | [109 127 79] | Internal capsule | |
| | | 2633 | 3.59 | <0.001 | 1.08 | [67 101 106] | Corona radiata | Right Superior |
| | | 1281 | 5.41 | <0.001 | 1.63 | [78 131 74] | Internal capsule | Right Anterior |
| | | 622 | 3.52 | <0.001 | 1.06 | [71 133 109] | Corona radiata | Right Superior |
| | | 10052 | -5.53 | <0.001 | 1.67 | [57 74 77] | Thalamic radiation | |
| | | 2085 | -4.13 | <0.001 | 1.24 | [67 70 39] | Cerebellar peduncle | Middle |
| | | 1063 | -4.84 | <0.001 | 1.46 | [126 67 83] | Thalamic radiation | |
| | | 828 | -5.17 | <0.001 | 1.56 | [89 84 83] | CC | Splenium |
| | | 694 | -4.28 | <0.001 | 1.29 | [96 88 50] | Cerebellar peduncle | Left Superior |
| | | 619 | -4.52 | <0.001 | 1.36 | [100 82 77] | CC | |

| | | | | | | | | |
|---|---|---|---|---|---|---|---|---|
| | ***Early Childhood Composite*** | 5784 | 3.83 | <0.001 | 1.15 | [114 117 96] | Corona radiata | Left Superior |
| | | 2734 | 3.98 | <0.001 | 1.20 | [56 127 94] | Longitudinal fasciculus | Right Superior |
| | | 1444 | 5.33 | <0.001 | 1.61 | [78 133 74] | Internal capsule | Right Anterior |
| | | 884 | 3.70 | <0.001 | 1.11 | [79 154 62] | CC | |
| | | 867 | 3.76 | <0.001 | 1.13 | [71 133 110] | | Right Superior |
| | | 701 | 4.04 | <0.001 | 1.22 | [106 95 41] | Cerebellar peduncle | Middle |
| | | 608 | 3.50 | 0.001 | 1.05 | [103 138 96] | CC | Body |
| | | 10772 | -6.15 | <0.001 | 1.85 | [50 84 82] | Longitudinal fasciculus | Right Superior |
| | | 1336 | -4.55 | <0.001 | 1.37 | [67 83 40] | Cerebellar peduncle | Middle |
| | | 1333 | -5.00 | <0.001 | 1.51 | [83 93 87] | CC | Splenium |
| | | 1129 | -4.04 | <0.001 | 1.22 | [126 67 84] | Thalamic radiation | |
| **ISO** | ***CCC*** | 7480 | -5.73 | <0.001 | 1.73 | [57 131 64] | External capsule | Right |
| | | 1493 | -4.20 | <0.001 | 1.26 | [123 117 56] | Fornix | Left |
| | | 936 | -4.25 | <0.001 | 1.28 | [94 89 91] | CC | Splenium |
| | | 613 | -3.96 | <0.001 | 1.19 | [75 113 71] | Internal capsule | Right Posterior |
| | | 587 | -3.72 | <0.001 | 1.12 | [75 78 96] | CC | Splenium |
| | | 563 | -4.90 | <0.001 | 1.48 | [81 131 97] | | Body |
| | | 423 | -3.66 | 0.008 | 1.10 | [123 64 90] | Thalamic radiation | |
| | | 422 | -4.06 | 0.008 | 1.22 | [128 96 103] | Longitudinal fasciculus | Left Superior |
| | | 422 | -3.31 | 0.008 | 1.00 | [57 108 108] | | Right Superior |
| | ***Fluid Composite*** | 10760 | -5.10 | <0.001 | 1.54 | [76 133 102] | CC | Body |
| | | 975 | -4.64 | <0.001 | 1.40 | [75 77 98] | | |
| | | 935 | -3.78 | <0.001 | 1.14 | [97 87 95] | CC | Splenium |
| | | 876 | -3.08 | <0.001 | 0.93 | [97 97 107] | | Left |
| | | 579 | -3.60 | 0.002 | 1.08 | [126 93 107] | Longitudinal fasciculus | Left Superior |
| | | 519 | -3.90 | 0.006 | 1.17 | [103 155 72] | CC | Genu |
| | | 504 | -3.05 | 0.008 | 0.92 | [55 64 71] | Thalamic radiation | Right Posterior |
| | | 471 | -2.76 | 0.013 | 0.83 | [111 118 110] | Corona radiata | Left Superior |
| | ***Early Childhood Composite*** | 447 | 5.03 | 0.009 | 1.51 | [100 108 48] | Cerebral peduncle | |
| | | 5544 | -4.80 | <0.001 | 1.45 | [76 133 102] | CC | Body |
| | | 665 | -4.84 | <0.001 | 1.46 | [86 101 99] | CC | Body |
| | | 437 | -4.33 | 0.011 | 1.30 | [127 93 107] | Longitudinal fasciculus | Left Superior |
| | | 428 | -4.03 | 0.013 | 1.21 | [108 81 100] | CC | Splenium |
| | | 407 | -3.40 | 0.019 | 1.02 | [61 105 73] | Internal capsule | Right Retrolenticular |
| **ODIt** | ***CCC*** | 7005 | -5.71 | <0.001 | 1.72 | [64 119 103] | Corona radiata | Right Superior |
| | | 2088 | -4.47 | <0.001 | 1.35 | [106 104 103] | CC | Body |

| | | | | | | | | |
|---|---|---|---|---|---|---|---|---|
| | | 758 | -4.87 | <0.001 | 1.47 | [114 127 102] | Corona radiata | Left Superior |
| | | 525 | -3.66 | <0.001 | 1.1 | [87 90 53] | Cerebellar peduncle | Right Superior |
| | | 461 | -4.12 | <0.001 | 1.24 | [96 84 41] | Cerebellar peduncle | Left Superior |
| | | 441 | -4.2 | <0.001 | 1.26 | [91 120 89] | Fornix | |
| | | 430 | -5.07 | <0.001 | 1.53 | [101 106 53] | Cerebral peduncle | Left |
| | | 385 | -4.7 | 0.001 | 1.42 | [63 91 92] | Corona radiata | Right Posterior |
| | | 379 | -4.23 | 0.001 | 1.27 | [106 134 105] | CC | Body |
| | | 332 | -4.95 | 0.005 | 1.49 | [119 77 85] | Tapetum | Left |
| | | 325 | -3.94 | 0.006 | 1.19 | [109 89 40] | Cerebellar peduncle | Middle |
| | | 315 | -4.44 | 0.008 | 1.34 | [118 84 97] | Corona radiata | Left Posterior |
| | | 309 | -4.99 | 0.009 | 1.5 | [128 90 101] | Longitudinal fasciculus | Left Superior |
| | | 300 | -4.32 | 0.012 | 1.3 | [59 108 74] | External capsule | Right |
| | | 277 | -4.47 | 0.024 | 1.35 | [55 80 76] | Thalamic radiation | Right Posterior |
| | ***Fluid Composite*** | 939 | 5.77 | <0.001 | 1.74 | [120 129 76] | External capsule | Left |
| | | 510 | 4.54 | <0.001 | 1.37 | [95 97 95] | CC | Body |
| | | 265 | 4.76 | 0.014 | 1.43 | [109 159 87] | | Left Anterior |
| | | 1752 | -5.81 | <0.001 | 1.75 | [64 120 101] | Corona radiata | Right Superior |
| | | 1225 | -5.35 | <0.001 | 1.61 | [79 90 80] | CC | Splenium |
| | | 936 | -5.42 | <0.001 | 1.63 | [115 100 96] | Corona radiata | Left Posterior |
| | | 569 | -4.65 | <0.001 | 1.4 | [105 105 56] | Cerebral peduncle | Left |
| | | 402 | -4.89 | <0.001 | 1.47 | [73 126 84] | Internal capsule | Right Anterior |
| | | 389 | -4.13 | <0.001 | 1.24 | [53 95 75] | | Right Retrolenticular |
| | | 310 | -4.16 | 0.003 | 1.25 | [110 107 105] | Corona radiata | Left Superior |
| | | 289 | -4.08 | 0.007 | 1.23 | [52 77 74] | Thalamic radiation | Right Posterior |
| | ***Early Childhood Composite*** | 1039 | 4.79 | <0.001 | 1.44 | [120 129 76] | External capsule | Left |
| | | 387 | 4.57 | <0.001 | 1.38 | [95 97 95] | CC | Body |
| | | 1152 | -5.42 | <0.001 | 1.63 | [118 92 101] | Corona radiata | Left Posterior |
| | | 1068 | -6.64 | <0.001 | 2 | [115 100 95] | | Left Posterior |
| | | 427 | -5.17 | <0.001 | 1.56 | [105 104 58] | Cerebral peduncle | Left |
| | | 405 | -3.16 | <0.001 | 0.95 | [75 150 91] | CC | Genu |
| | | 386 | -4.37 | <0.001 | 1.32 | [109 107 105] | Corona radiata | Left Superior |
| | | 307 | -4.69 | 0.005 | 1.41 | [124 97 99] | Longitudinal fasciculus | Left Superior |
| | | 262 | -4.14 | 0.021 | 1.25 | [58 58 78] | Thalamic radiation | Right Posterior |
| **ODIp** | ***CCC*** | 331 | 4.78 | 0.008 | 1.44 | [125 116 75] | External capsule | Left |
| | | 2261 | -5.32 | <0.001 | 1.6 | [64 107 69] | Internal capsule | Right Retrolenticular |
| | | 2152 | -4.43 | <0.001 | 1.33 | [77 150 86] | CC | Genu |
| | | 1543 | -4.99 | <0.001 | 1.5 | [120 83 95] | Corona radiata | Left Posterior |

| | | | | | | | | |
|---|---|---|---|---|---|---|---|---|
| | | 1319 | -4.73 | <0.001 | 1.42 | [74 121 110] | CC | Body |
| | | 1087 | -3.94 | <0.001 | 1.19 | [114 105 87] | Internal capsule | Left Posterior |
| | | 665 | -4.82 | <0.001 | 1.45 | [83 84 42] | Cerebellar peduncle | Right Superior |
| | | 654 | -4.01 | <0.001 | 1.21 | [98 89 40] | Medial lemniscus | Left |
| | | 322 | -5.37 | 0.011 | 1.62 | [104 129 78] | Internal capsule | Left Anterior |
| | | 319 | -3.78 | 0.012 | 1.14 | [110 89 40] | Cerebellar peduncle | Middle |
| | ***Fluid Composite*** | 430 | 5.61 | <0.001 | 1.69 | [120 129 76] | External capsule | Left |
| | | 2301 | -5.38 | <0.001 | 1.62 | [101 87 79] | CC | Splenium |
| | | 1319 | -4.41 | <0.001 | 1.33 | [72 113 72] | Internal capsule | Right Posterior |
| | | 930 | -5.11 | <0.001 | 1.54 | [116 104 97] | Corona radiata | Left Superior |
| | | 702 | -5.29 | <0.001 | 1.59 | [77 153 76] | CC | Genu |
| | | 524 | -5.71 | <0.001 | 1.72 | [105 106 54] | Cerebral peduncle | Left |
| | | 483 | -4.67 | <0.001 | 1.41 | [76 128 79] | Internal capsule | Right Anterior |
| | | 430 | -5.14 | <0.001 | 1.55 | [87 92 55] | Cerebellar peduncle | Right Superior |
| | | 420 | -5.99 | <0.001 | 1.8 | [75 79 39] | Cerebellar peduncle | Middle |
| | | 358 | -5.24 | 0.003 | 1.58 | [99 90 45] | Medial lemniscus | Left |
| | | 355 | -4.92 | 0.003 | 1.48 | [84 153 72] | CC | Genu |
| | | 327 | -3.79 | 0.006 | 1.14 | [91 118 86] | Fornix | |
| | | 325 | -4.63 | 0.006 | 1.39 | [52 95 70] | Internal capsule | Right Retrolenticular |
| | | 286 | -4.1 | 0.019 | 1.23 | [61 53 74] | Thalamic radiation | Right Posterior |
| | ***Early Childhood Composite*** | 433 | 4.78 | <0.001 | 1.44 | [120 129 76] | External capsule | Left |
| | | 2645 | -5.23 | <0.001 | 1.58 | [72 115 109] | Corona radiata | Right Superior |
| | | 2587 | -6.19 | <0.001 | 1.86 | [110 76 98] | | Left Posterior |
| | | 1532 | -5.49 | <0.001 | 1.65 | [72 160 73] | | Right Anterior |
| | | 950 | -5.82 | <0.001 | 1.75 | [114 102 92] | | Left Posterior |
| | | 729 | -5 | <0.001 | 1.51 | [54 74 73] | Thalamic radiation | Right Posterior |
| | | 645 | -5.26 | <0.001 | 1.58 | [84 84 45] | Cerebellar peduncle | Right Superior |
| | | 446 | -4.52 | <0.001 | 1.36 | [105 104 58] | Cerebral peduncle | Left |
| | | 373 | -4.99 | 0.003 | 1.5 | [99 90 45] | Medial lemniscus | Left |
| | | 4079 | -5.87 | <0.001 | 1.77 | [51 105 103] | Longitudinal fasciculus | Right Superior |
| **ODIs** | ***CCC*** | 45066 | -6.10 | <0.001 | 1.84 | [69 158 87] | Corona radiata | Right Anterior |
| | | 2495 | -5.97 | <0.001 | 1.80 | [128 102 107] | Longitudinal fasciculus | Left Superior |
| | ***Fluid Composite*** | 27011 | -5.83 | <0.001 | 1.76 | [97 89 81] | CC | Splenium |
| | | 1144 | -4.65 | <0.001 | 1.40 | [103 92 31] | Cerebellar peduncle | Middle |
| | | 1040 | -3.84 | <0.001 | 1.16 | [128 115 101] | Longitudinal fasciculus | Left Superior |
| | | 966 | -5.31 | <0.001 | 1.60 | [69 83 33] | Cerebellar peduncle | Middle |
| | | 637 | -4.18 | 0.003 | 1.26 | [95 80 45] | | Left Superior |

| | | | | | | | |
|---|---|---|---|---|---|---|---|
| ***Early Childhood Composite*** | 33142 | -6.09 | <0.001 | 1.83 | [80 146 66] | CC | Genu |
| | 1523 | -3.88 | <0.001 | 1.17 | [54 101 104] | Longitudinal fasciculus | Right Superior |
| | 1457 | -4.00 | <0.001 | 1.20 | [122 110 110] | | Left Superior |
| | 1164 | -4.73 | <0.001 | 1.42 | [103 92 31] | Cerebellar peduncle | Middle |
| | 721 | -4.38 | 0.001 | 1.32 | [94 95 36] | | Middle |
| | 591 | -5.81 | 0.007 | 1.75 | [105 103 55] | Cerebral peduncle | |
| | 584 | -3.85 | 0.008 | 1.16 | [102 127 99] | CC | Body |

Grey cells indicate a positive t-statistic and white cells indicate a negative t-statistic. Significance was determined using a permutation cluster-wise $P < 0.050$ familywise error (FWE)-corrected threshold. Positive t-statistic values indicate a significantly positive regression between the diffusion signal and cognitive test, while negative t-statistic values indicate a significantly negative regression in the HC group (HC n = 21). *Abbreviations: [X Y Z], coordinates of the voxels on the MNI 1 mm brain atlas; CC, corpus callosum; CCC, Cognitive Crystallized Composite; CS, cancer survivors; FWE, familywise error; HC, healthy controls; HCP, Human Connectome Project; ISO, isotropic volume fraction; NDI, neurite density index; ODIp, primary orientation dispersion index; ODIs, secondary orientation dispersion index; ODIt, total orientation dispersion index; t, t-statistic; WM, white matter.*

**Supplementary Table 3** Significant voxel cluster associations between NODDI-Bingham metrics and cognitive scores in the CS group at the $P < 0.05$ FWE-corrected level.

| NODDI metric | Cognitive test | Voxels | Peak t-stat | P-Value | Peak Cohen's d | Coordinates [X Y Z] | Region | Subregion |
|---|---|---|---|---|---|---|---|---|
| **NDI** | ***CCC*** | 1380 | -4.80 | <0.001 | 1.45 | [113 159 73] | Corona radiata | Left Anterior |
| | | 887 | -3.84 | <0.001 | 1.16 | [125 83 103] | Longitudinal fasciculus | |
| | | 886 | -4.32 | <0.001 | 1.30 | [58 115 64] | External capsule | Right |
| | | 877 | -3.53 | <0.001 | 1.06 | [67 116 87] | Internal capsule | Right Posterior |
| | | 622 | -3.49 | <0.001 | 1.05 | [111 149 99] | Corona radiata | Left Anterior |
| | ***Fluid Composite*** | 3601 | 4.18 | <0.001 | 1.26 | [78 80 38] | Cerebellar peduncle | Right Inferior |
| | | 587 | 3.83 | <0.001 | 1.15 | [87 102 95] | CC | Body |
| | | 531 | 4.67 | <0.001 | 1.41 | [76 80 74] | | |
| | | 364 | 3.31 | 0.024 | 1.00 | [81 77 96] | | |
| | ***Early Childhood Composite*** | 4402 | 4.89 | <0.001 | 1.47 | [93 91 39] | Medial lemniscus | Left |
| | | 839 | 4.54 | <0.001 | 1.37 | [88 101 95] | CC | Body |
| | | 620 | 5.16 | <0.001 | 1.55 | [76 79 75] | | |
| **ISO** | ***CCC*** | 549 | -3.21 | <0.001 | 0.97 | [118 131 100] | Internal capsule | |
| | | 312 | -3.31 | 0.004 | 1.00 | [51 91 63] | Sagittal stratum | Right |
| | | 305 | -3.40 | 0.005 | 1.02 | [97 110 42] | Cerebellar peduncle | Middle |
| | ***Fluid Composite*** | 1770 | -4.97 | <0.001 | 1.50 | [74 80 36] | Cerebellar peduncle | Middle |
| | | 454 | -3.87 | <0.001 | 1.17 | [117 73 36] | | Middle |
| | | 361 | -4.12 | 0.008 | 1.24 | [123 110 82] | External capsule | Left |
| | | 346 | -4.50 | 0.011 | 1.36 | [101 79 40] | Cerebellar peduncle | Left Inferior |
| | ***Early Childhood Composite*** | 1187 | -4.07 | <0.001 | 1.23 | [71 78 37] | Cerebellar peduncle | Middle |
| | | 463 | -4.25 | <0.001 | 1.28 | [123 109 83] | Internal capsule | |
| | | 412 | -4.83 | 0.001 | 1.45 | [116 99 79] | | Left Retrolenticular |
| | | 348 | -4.60 | 0.006 | 1.39 | [134 99 63] | External capsule | |
| | | 318 | -3.55 | 0.014 | 1.07 | [101 79 39] | Cerebellar peduncle | Left Inferior |
| | | 305 | -3.41 | 0.019 | 1.03 | [61 135 73] | External capsule | Right |
| **ODIt** | ***CCC*** | 443 | -4.21 | <0.001 | 1.27 | [119 100 98] | Corona radiata | |
| | | 295 | -4.33 | 0.002 | 1.30 | [83 112 102] | CC | Body |
| | | 287 | -4.47 | 0.003 | 1.35 | [101 144 94] | | Body |
| | | 271 | -4.04 | 0.005 | 1.22 | [102 125 103] | | Body |
| | ***Fluid Composite*** | 564 | -4.64 | <0.001 | 1.40 | [113 160 78] | Corona radiata | Left Anterior |
| | | 457 | -4.20 | <0.001 | 1.26 | [101 155 69] | | Genu |

| | | | | | | | | |
|---|---|---|---|---|---|---|---|---|
| | | 456 | -4.36 | <0.001 | 1.31 | [74 92 102] | | Splenium |
| | | 404 | -4.00 | <0.001 | 1.20 | [125 85 78] | Thalamic radiation | Left Posterior |
| | | 340 | -5.40 | 0.001 | 1.63 | [103 151 92] | CC | Genu |
| | | 270 | -4.60 | 0.012 | 1.39 | [80 159 80] | Cingulum | |
| | | 257 | -4.09 | 0.019 | 1.23 | [108 93 100] | CC | Splenium |
| | ***Early Childhood Composite*** | 8495 | -7.50 | <0.001 | 2.26 | [80 159 80] | Cingulum | |
| | | 520 | -4.14 | <0.001 | 1.25 | [65 101 95] | Corona radiata | Right Posterior |
| | | 495 | -5.85 | <0.001 | 1.76 | [125 85 79] | Thalamic radiation | Left Posterior |
| | | 495 | -4.80 | <0.001 | 1.45 | [102 89 95] | CC | Splenium |
| | | 421 | -5.56 | 0.002 | 1.67 | [124 92 104] | Longitudinal fasciculus | Left Superior |
| **ODIp** | ***CCC*** | 333 | -4.29 | <0.001 | 1.29 | [102 126 100] | CC | Body |
| | | 311 | -4.68 | 0.001 | 1.41 | [101 141 96] | | Body |
| | | 288 | -4.78 | 0.003 | 1.44 | [110 99 106] | | Left Posterior |
| | | 262 | -4.93 | 0.007 | 1.48 | [73 100 39] | Cerebellar peduncle | Middle |
| | ***Fluid Composite*** | 289 | -4.72 | 0.002 | 1.42 | [75 92 102] | CC | Splenium |
| | | 276 | -5.34 | 0.003 | 1.61 | [111 142 84] | | Left Anterior |
| | | 221 | -4.64 | 0.022 | 1.40 | [101 90 99] | | Splenium |
| | | 219 | -4.65 | 0.023 | 1.40 | [107 153 96] | | Left Anterior |
| | ***Early Childhood Composite*** | 1975 | -7.71 | <0.001 | 2.32 | [75 92 102] | CC | Splenium |
| | | 1114 | -4.36 | <0.001 | 1.31 | [107 108 105] | | Body |
| | | 250 | -4.49 | 0.020 | 1.35 | [102 89 95] | | Splenium |
| **ODIs** | ***CCC*** | 489 | -4.67 | <0.001 | 1.41 | [76 113 103] | CC | Body |
| | | 327 | -3.08 | 0.008 | 0.93 | [115 102 104] | Corona radiata | Left Superior |
| | ***Fluid Composite*** | 2002 | -4.40 | <0.001 | 1.33 | [102 154 90] | CC | Genu |
| | | 510 | -3.32 | 0.001 | 1.00 | [107 76 41] | Cerebellar peduncle | Middle |
| | | 491 | -5.17 | 0.002 | 1.56 | [122 85 105] | Longitudinal fasciculus | Left Superior |
| | | 434 | -4.09 | 0.005 | 1.23 | [127 96 70] | Internal capsule | Left Retrolenticular |
| | | 361 | -3.87 | 0.024 | 1.17 | [125 69 70] | Thalamic radiation | Left Posterior |
| | ***Early Childhood Composite*** | 3648 | -5.12 | <0.001 | 1.54 | [93 154 80] | CC | Genu |
| | | 1446 | -4.80 | <0.001 | 1.45 | [105 88 103] | | Splenium |
| | | 1105 | -7.78 | <0.001 | 2.34 | [122 85 105] | Longitudinal fasciculus | Left Superior |
| | | 1081 | -4.13 | <0.001 | 1.24 | [109 72 40] | Cerebellar peduncle | Middle |
| | | 714 | -4.61 | <0.001 | 1.39 | [126 102 66] | Sagittal stratum | Left |
| | | 545 | -3.93 | 0.002 | 1.18 | [74 79 40] | Cerebellar peduncle | Middle |
| | | 501 | -4.84 | 0.004 | 1.46 | [76 74 95] | CC | |

| | | | | | | | | |
|---|---|---|---|---|---|---|---|---|
| | | 475 | -4.68 | 0.007 | 1.41 | [126 80 87] | Longitudinal fasciculus | Left Superior |

Grey cells indicate a positive t-statistic and white cells indicate a negative t-statistic. Significance was determined using a permutation cluster-wise *P* < 0.050 familywise error (FWE)-corrected threshold. Positive t-statistic values indicate a significantly positive regression between the diffusion signal and cognitive test, while negative t-statistic values indicate a significantly negative regression in the CS group (CS n = 21). *Abbreviations: [X Y Z], coordinates of the voxels on the MNI 1 mm brain atlas; CC, corpus callosum; CCC, Cognitive Crystallized Composite; CS, cancer survivors; FWE, familywise error; HC, healthy controls; HCP, Human Connectome Project; ISO, isotropic volume fraction; NDI, neurite density index; ODIp, primary orientation dispersion index; ODIs, secondary orientation dispersion index; ODIt, total orientation dispersion index; t, t-statistic; WM, white matter.*

**Supplementary Table 4** Significant voxel cluster associations between NODDI-Bingham metrics and cognitive scores in the HCP control group ages 22–30 at the P < 0.05 FWE-corrected level.

| NODDI metric | Cognitive test | Voxels | Peak t-stat | P-Value | Peak Cohen's d | Coordinates [X Y Z] | Region | Subregion |
|---|---|---|---|---|---|---|---|---|
| **NDI** | CCC | 294 | 3.13 | 0.020 | 0.12 | [112 119 80] | Internal capsule | |
| | | 4285 | -5.39 | <0.001 | 0.21 | [62 95 77] | | |
| | | 4221 | -4.90 | <0.001 | 0.19 | [84 93 41] | | |
| | | 352 | -4.85 | 0.004 | 0.19 | [58 88 98] | | |
| | | 2095 | -4.65 | <0.001 | 0.18 | [73 82 80] | CC | Splenium |
| | | 1477 | -4.69 | <0.001 | 0.18 | [94 140 90] | | Body |
| | | 749 | -3.41 | <0.001 | 0.13 | [75 163 78] | | Genu |
| | | 704 | -3.76 | <0.001 | 0.14 | [72 113 105] | | Body |
| | | 620 | -3.82 | <0.001 | 0.15 | [100 82 93] | | Splenium |
| | | 1124 | -4.34 | <0.001 | 0.17 | [74 83 32] | Cerebellar peduncle | Middle |
| | | 1057 | -4.56 | <0.001 | 0.17 | [121 79 87] | Thalamic radiation | Left Posterior |
| | | 375 | -4.48 | 0.002 | 0.17 | [128 91 67] | Sagittal stratum | Left |
| | Fluid Composite | 257 | 4.70 | 0.002 | 0.18 | [59 103 72] | Internal capsule | Right Retrolenticular |
| | | 253 | 3.63 | 0.002 | 0.14 | [131 112 100] | Longitudinal fasciculus | Left Superior |
| | | 222 | 3.31 | 0.008 | 0.13 | [123 91 104] | | Left Superior |
| | | 234 | 4.83 | 0.005 | 0.19 | [63 70 93] | Corona radiata | Right Posterior |
| | | 209 | 3.37 | 0.013 | 0.13 | [103 123 106] | CC | Body |
| | | 535 | -4.79 | <0.001 | 0.18 | [70 70 85] | Thalamic radiation | |
| | | 512 | -3.90 | <0.001 | 0.15 | [109 64 40] | | |

| NODDI metric | Cognitive test | Voxels | Peak t-stat | P-Value | Peak Cohen's d | Coordinates [X Y Z] | Region | Subregion |
|---|---|---|---|---|---|---|---|---|
| | | 383 | -3.65 | <0.001 | 0.14 | [83 80 86] | | |
| | | 230 | -3.61 | 0.005 | 0.14 | [80 99 36] | | |
| | | 201 | -4.51 | 0.018 | 0.17 | [55 109 62] | | |
| | | 287 | -3.54 | <0.001 | 0.14 | [60 93 78] | Internal capsule | Right Retrolenticular |
| | | 272 | -3.20 | 0.001 | 0.12 | [96 116 102] | CC | Body |
| | | 221 | -3.88 | 0.008 | 0.15 | [80 122 103] | | Body |
| | Early Childhood Composite | 261 | 4.61 | 0.004 | 0.18 | [59 103 72] | Internal capsule | Right Retrolenticular |
| | | 1712 | -4.38 | <0.001 | 0.17 | [111 66 40] | Cerebellar peduncle | Middle |
| | | 333 | -3.72 | <0.001 | 0.14 | [72 76 39] | | Middle |
| | | 955 | -4.13 | <0.001 | 0.16 | [100 81 92] | CC | Splenium |
| | | 435 | -3.66 | <0.001 | 0.14 | [81 81 87] | | Splenium |
| | | 361 | -4.69 | <0.001 | 0.18 | [80 112 103] | | Body |
| | | 764 | -3.99 | <0.001 | 0.15 | [60 92 80] | Internal capsule | |
| | | 337 | -4.34 | <0.001 | 0.17 | [70 70 85] | | |
| | | 274 | -3.93 | 0.002 | 0.15 | [59 116 63] | | |
| | | 216 | -3.80 | 0.021 | 0.15 | [123 121 86] | | |
| | | 555 | -5.33 | <0.001 | 0.20 | [79 103 47] | Corticospinal tract | Right |
| | | 231 | -4.19 | 0.012 | 0.16 | [121 80 86] | Thalamic radiation | Left Posterior |
| **ISO** | CCC | 2990 | 4.06 | <0.001 | 0.16 | [111 134 108] | Corona radiata | Left Superior |
| | | 1117 | 4.20 | <0.001 | 0.16 | [67 117 82] | Internal capsule | Right Posterior |
| | | 880 | 4.93 | <0.001 | 0.19 | [65 93 88] | | Right Retrolenticular |
| | | 418 | 4.01 | <0.001 | 0.15 | [85 142 92] | CC | Body |
| | | 356 | 3.92 | 0.001 | 0.15 | [92 117 100] | | Body |
| | | 365 | 3.54 | <0.001 | 0.14 | [114 91 93] | Sagittal stratum | |
| | | 282 | 4.38 | 0.010 | 0.17 | [77 90 30] | Cerebellar peduncle | Middle |
| | | 1280 | -4.17 | <0.001 | 0.16 | [97 89 84] | CC | Splenium |
| | Fluid Composite | 725 | 3.69 | <0.001 | 0.14 | [65 93 79] | Fornix | Right |
| | | 608 | 4.62 | <0.001 | 0.18 | [104 95 32] | Cerebellar peduncle | Middle |
| | | 217 | 4.20 | 0.022 | 0.16 | [80 101 29] | | Middle |
| | | 529 | 3.91 | <0.001 | 0.15 | [94 142 92] | CC | Body |
| | | 487 | 3.35 | <0.001 | 0.13 | [102 129 78] | Internal capsule | Left Anterior |
| | | 1752 | -5.20 | <0.001 | 0.20 | [86 91 81] | CC | Splenium |

| **NODDI metric** | **Cognitive test** | **Voxels** | **Peak t-stat** | **P-Value** | **Peak Cohen's d** | **Coordinates [X Y Z]** | **Region** | **Subregion** |
|---|---|---|---|---|---|---|---|---|
| | | 441 | -4.17 | <0.001 | 0.16 | [86 98 29] | Corticospinal tract | Right |
| | | 231 | -3.36 | 0.013 | 0.13 | [101 102 50] | | Left |
| | | 303 | -4.41 | <0.001 | 0.17 | [111 66 40] | Cerebellar peduncle | Middle |
| | | 282 | -4.03 | 0.002 | 0.15 | [49 90 59] | Sagittal stratum | Right |
| | | 227 | -4.46 | 0.015 | 0.17 | [97 96 33] | Cerebellar peduncle | |
| | Early Childhood Composite | 1012 | 5.09 | <0.001 | 0.20 | [107 95 33] | Cerebellar peduncle | Middle |
| | | 244 | 4.18 | 0.017 | 0.16 | [80 101 29] | | Middle |
| | | 527 | 3.98 | <0.001 | 0.15 | [66 93 80] | Fornix | Right |
| | | 1983 | -4.45 | <0.001 | 0.17 | [86 91 81] | CC | Splenium |
| | | 1542 | -5.27 | <0.001 | 0.20 | [111 66 40] | Cerebellar peduncle | Middle |
| | | 700 | -4.52 | <0.001 | 0.17 | [80 87 33] | | Right Inferior |
| | | 441 | -3.57 | <0.001 | 0.14 | [96 100 43] | Corticospinal tract | Left |
| **ODIt** | CCC | 971 | 4.47 | <0.001 | 0.17 | [102 159 81] | CC | Genu |
| | | 408 | 4.22 | <0.001 | 0.16 | [79 160 73] | | Genu |
| | | 285 | 3.78 | <0.001 | 0.15 | [71 109 106] | | Body |
| | | 184 | 3.58 | 0.022 | 0.14 | [101 82 93] | | Splenium |
| | | 698 | 4.72 | <0.001 | 0.18 | [66 100 79] | Internal capsule | Right Retrolenticular |
| | | 485 | 4.37 | <0.001 | 0.17 | [104 115 70] | | Left Posterior |
| | | 293 | 4.43 | <0.001 | 0.17 | [72 113 70] | | Right Posterior |
| | | 676 | 5.54 | <0.001 | 0.21 | [80 80 76] | CC | |
| | | 201 | 4.52 | 0.010 | 0.17 | [61 83 82] | | |
| | | 469 | 4.21 | <0.001 | 0.16 | [68 154 81] | Corona radiata | Right Anterior |
| | | 1627 | -4.82 | <0.001 | 0.18 | [92 91 84] | CC | Splenium |
| | | 289 | -3.84 | <0.001 | 0.15 | [87 124 98] | | Body |
| | | 794 | -4.90 | <0.001 | 0.19 | [128 81 72] | Thalamic radiation | Left Posterior |
| | | 735 | -4.65 | <0.001 | 0.18 | [66 82 104] | Corona radiata | Right Posterior |
| | | 241 | -3.86 | 0.002 | 0.15 | [114 90 106] | | Left Posterior |
| | | 485 | -4.85 | <0.001 | 0.19 | [123 98 108] | Longitudinal fasciculus | Left Superior |
| | | 332 | -4.30 | <0.001 | 0.17 | [56 97 105] | | Right Superior |
| | | 298 | -5.08 | <0.001 | 0.19 | [59 112 107] | | Right Superior |
| | | 454 | -4.07 | <0.001 | 0.16 | [114 82 35] | Cerebellar peduncle | Middle |
| | | 349 | -5.04 | <0.001 | 0.19 | [85 90 51] | | Right Superior |

| NODDI metric | Cognitive test | Voxels | Peak t-stat | P-Value | Peak Cohen's d | Coordinates [X Y Z] | Region | Subregion |
|---|---|---|---|---|---|---|---|---|
| | | 432 | -3.69 | <0.001 | 0.14 | [119 106 90] | External capsule | Left |
| | | 293 | -4.54 | <0.001 | 0.17 | [61 111 87] | | Right |
| | | 208 | -3.73 | 0.008 | 0.14 | [80 95 35] | Cerebellar peduncle | |
| | | 182 | -3.96 | 0.024 | 0.15 | [73 108 68] | Internal capsule | Right Posterior |
| | Fluid Composite | 298 | 3.75 | <0.001 | 0.14 | [67 146 70] | External capsule | Right |
| | | 298 | 4.14 | <0.001 | 0.16 | [60 83 82] | Tapetum | |
| | | 264 | 3.91 | <0.001 | 0.15 | [85 81 89] | CC | |
| | | 254 | 4.36 | <0.001 | 0.17 | [68 68 81] | | |
| | | 214 | 3.97 | 0.005 | 0.15 | [52 88 61] | Sagittal stratum | Right |
| | | 188 | 4.11 | 0.016 | 0.16 | [65 103 72] | Internal capsule | Right Retrolenticular |
| | | 3838 | -4.77 | <0.001 | 0.18 | [90 91 81] | CC | Splenium |
| | | 1457 | -5.12 | <0.001 | 0.20 | [68 73 34] | Cerebellar peduncle | Middle |
| | | 531 | -4.43 | <0.001 | 0.17 | [112 81 33] | | Middle |
| | | 189 | -3.96 | 0.015 | 0.15 | [112 62 34] | | Middle |
| | | 787 | -4.55 | <0.001 | 0.17 | [118 122 96] | Corona radiata | Left Superior |
| | | 465 | -4.72 | <0.001 | 0.18 | [61 87 92] | | Right Posterior |
| | | 310 | -4.92 | <0.001 | 0.19 | [121 60 71] | Thalamic radiation | Left Posterior |
| | | 305 | -4.24 | <0.001 | 0.16 | [60 64 74] | | Right Posterior |
| | | 214 | -3.46 | 0.005 | 0.13 | [122 79 84] | | Left Posterior |
| | | 225 | -4.18 | 0.003 | 0.16 | [125 81 99] | Longitudinal fasciculus | |
| | Early Childhood Composite | 434 | 4.54 | <0.001 | 0.17 | [66 100 79] | Internal capsule | Right Retrolenticular |
| | | 414 | 3.81 | <0.001 | 0.15 | [70 152 85] | Corona radiata | Right Anterior |
| | | 385 | 3.73 | <0.001 | 0.14 | [107 164 83] | | Left Anterior |
| | | 323 | 3.75 | <0.001 | 0.14 | [51 105 101] | Longitudinal fasciculus | Right Superior |
| | | 312 | 4.16 | <0.001 | 0.16 | [78 87 99] | CC | Splenium |
| | | 281 | 3.71 | <0.001 | 0.14 | [97 136 98] | | Body |
| | | 204 | 4.06 | 0.011 | 0.16 | [60 83 82] | Tapetum | |
| | | 189 | 4.04 | 0.020 | 0.16 | [67 66 81] | Thalamic radiation | |
| | | 4374 | -4.43 | <0.001 | 0.17 | [74 85 80] | CC | Splenium |
| | | 1213 | -4.94 | <0.001 | 0.19 | [80 98 32] | Cerebellar peduncle | Middle |

| NODDI metric | Cognitive test | Voxels | Peak t-stat | P-Value | Peak Cohen's d | Coordinates [X Y Z] | Region | Subregion |
|---|---|---|---|---|---|---|---|---|
| | | 402 | -4.13 | <0.001 | 0.16 | [112 81 33] | | Middle |
| | | 302 | -4.30 | <0.001 | 0.17 | [113 66 38] | | Middle |
| | | 243 | -4.67 | 0.002 | 0.18 | [95 94 36] | | Middle |
| | | 811 | -5.32 | <0.001 | 0.20 | [118 122 96] | Corona radiata | Left Superior |
| | | 337 | -4.22 | <0.001 | 0.16 | [62 124 94] | | Right Superior |
| | | 512 | -4.68 | <0.001 | 0.18 | [125 81 99] | Longitudinal fasciculus | |
| | | 479 | -4.41 | <0.001 | 0.17 | [57 81 85] | Thalamic radiation | Right Posterior |
| | | 290 | -3.92 | <0.001 | 0.15 | [123 68 70] | | Left Posterior |
| | | 218 | -3.66 | 0.006 | 0.14 | [56 68 67] | Sagittal stratum | Right |
| | | 204 | -4.38 | 0.011 | 0.17 | [61 105 75] | Internal capsule | Right Retrolenticular |
| | | 200 | -4.59 | 0.013 | 0.18 | [54 97 104] | Longitudinal fasciculus | Right Superior |
| **ODIp** | CCC | 1142 | 5.03 | <0.001 | 0.19 | [58 96 74] | Internal capsule | Right Retrolenticular |
| | | 502 | 4.05 | <0.001 | 0.16 | [104 115 70] | | Left Posterior |
| | | 354 | 4.67 | <0.001 | 0.18 | [72 113 70] | | Right Posterior |
| | | 822 | 4.21 | <0.001 | 0.16 | [107 154 83] | CC | |
| | | 697 | 5.47 | <0.001 | 0.21 | [81 79 78] | | |
| | | 522 | 3.87 | <0.001 | 0.15 | [85 153 72] | CC | Genu |
| | | 393 | 3.67 | <0.001 | 0.14 | [69 158 77] | Corona radiata | Right Anterior |
| | | 345 | 3.54 | <0.001 | 0.14 | [73 117 109] | | Right Superior |
| | | 1637 | -4.67 | <0.001 | 0.18 | [92 91 83] | CC | Splenium |
| | | 362 | -3.99 | <0.001 | 0.15 | [77 123 102] | | Body |
| | | 1120 | -4.55 | <0.001 | 0.17 | [60 87 95] | Corona radiata | Right Posterior |
| | | 313 | -3.59 | <0.001 | 0.14 | [122 79 94] | | Left Posterior |
| | | 1042 | -4.29 | <0.001 | 0.16 | [122 103 72] | Internal capsule | Left Retrolenticular |
| | | 535 | -3.91 | <0.001 | 0.15 | [67 111 71] | | Right Posterior |
| | | 842 | -4.52 | <0.001 | 0.17 | [126 77 67] | Sagittal stratum | Left |
| | | 650 | -4.92 | <0.001 | 0.19 | [125 89 103] | Longitudinal fasciculus | Left Superior |
| | | 351 | -5.01 | <0.001 | 0.19 | [57 93 104] | | Right Superior |
| | | 264 | -4.73 | 0.004 | 0.18 | [58 111 106] | | Right Superior |
| | | 577 | -4.32 | <0.001 | 0.17 | [117 81 36] | Cerebellar peduncle | Middle |
| | | 371 | -5.43 | <0.001 | 0.21 | [86 91 53] | | Right Superior |
| | | 361 | -4.51 | <0.001 | 0.17 | [61 111 87] | External capsule | Right |

| NODDI metric | Cognitive test | Voxels | Peak t-stat | P-Value | Peak Cohen's d | Coordinates [X Y Z] | Region | Subregion |
|---|---|---|---|---|---|---|---|---|
| | | 244 | -3.49 | 0.008 | 0.13 | [69 137 95] | Superior fronto-occipital fasciculus | |
| | Fluid Composite | 408 | 4.87 | <0.001 | 0.19 | [58 96 74] | Internal capsule | Right Retrolenticular |
| | | 285 | 3.89 | 0.002 | 0.15 | [65 102 76] | | Right Retrolenticular |
| | | 403 | 4.96 | <0.001 | 0.19 | [78 87 99] | CC | Splenium |
| | | 361 | 4.46 | <0.001 | 0.17 | [68 68 81] | Thalamic radiation | |
| | | 215 | 3.72 | 0.024 | 0.14 | [68 146 70] | | |
| | | 225 | 4.27 | 0.017 | 0.16 | [59 79 82] | Tapetum | |
| | | 3892 | -4.75 | <0.001 | 0.18 | [90 91 81] | CC | Splenium |
| | | 1635 | -5.46 | <0.001 | 0.21 | [68 73 34] | Cerebellar peduncle | Middle |
| | | 987 | -4.36 | <0.001 | 0.17 | [118 109 90] | External capsule | Left |
| | | 355 | -3.59 | <0.001 | 0.14 | [60 112 84] | | Right |
| | | 840 | -4.36 | <0.001 | 0.17 | [115 63 33] | Cerebellar peduncle | |
| | | 316 | -4.54 | <0.001 | 0.17 | [125 81 99] | | |
| | | 506 | -4.93 | <0.001 | 0.19 | [121 60 71] | Thalamic radiation | Left Posterior |
| | | 414 | -3.96 | <0.001 | 0.15 | [58 67 73] | | Right Posterior |
| | | 499 | -5.15 | <0.001 | 0.20 | [59 81 91] | Corona radiata | Right Posterior |
| | | 219 | -4.14 | 0.021 | 0.16 | [82 86 39] | Medial lemniscus | Right |
| | Early Childhood Composite | 826 | 5.07 | <0.001 | 0.19 | [58 96 74] | Internal capsule | Right Retrolenticular |
| | | 379 | 3.49 | <0.001 | 0.13 | [115 163 80] | Cingulum | |
| | | 310 | 3.47 | <0.001 | 0.13 | [68 147 69] | | |
| | | 340 | 4.60 | <0.001 | 0.18 | [52 106 101] | Longitudinal fasciculus | Right Superior |
| | | 272 | 3.60 | 0.003 | 0.14 | [95 126 101] | CC | Body |
| | | 4087 | -4.39 | <0.001 | 0.17 | [74 85 80] | | Splenium |
| | | 1478 | -4.97 | <0.001 | 0.19 | [81 101 32] | Cerebellar peduncle | Middle |
| | | 421 | -4.37 | <0.001 | 0.17 | [111 86 37] | | Middle |
| | | 378 | -4.49 | <0.001 | 0.17 | [118 68 36] | | Middle |
| | | 358 | -3.58 | <0.001 | 0.14 | [79 86 35] | | Right Inferior |
| | | 255 | -4.94 | 0.006 | 0.19 | [95 94 36] | | Middle |
| | | 1088 | -5.52 | <0.001 | 0.21 | [120 109 84] | External capsule | Left |
| | | 518 | -4.49 | <0.001 | 0.17 | [60 112 84] | | Right |

| NODDI metric | Cognitive test | Voxels | Peak t-stat | P-Value | Peak Cohen's d | Coordinates [X Y Z] | Region | Subregion |
|---|---|---|---|---|---|---|---|---|
| | | 731 | -4.89 | <0.001 | 0.19 | [59 82 91] | Corona radiata | Right Posterior |
| | | 682 | -5.10 | <0.001 | 0.20 | [125 81 99] | Longitudinal fasciculus | |
| | | 500 | -3.69 | <0.001 | 0.14 | [126 73 70] | Thalamic radiation | Left Posterior |
| | | 262 | -3.38 | 0.005 | 0.13 | [58 67 73] | | Right Posterior |
| | | 262 | -4.38 | 0.005 | 0.17 | [62 105 78] | Internal capsule | Right Retrolenticular |
| | | 231 | -4.36 | 0.015 | 0.17 | [52 118 101] | Longitudinal fasciculus | Right Superior |
| **ODIs** | CCC | 2382 | 4.66 | <0.001 | 0.18 | [73 149 91] | CC | Genu |
| | | 399 | 4.25 | <0.001 | 0.16 | [71 108 105] | | Body |
| | | 298 | 3.74 | 0.008 | 0.14 | [98 83 93] | | Splenium |
| | | 668 | 4.54 | <0.001 | 0.17 | [63 93 83] | Internal capsule | |
| | | 468 | 4.32 | <0.001 | 0.17 | [81 80 77] | | |
| | | 378 | 4.23 | <0.001 | 0.16 | [117 114 108] | | |
| | | 379 | 4.03 | <0.001 | 0.15 | [76 117 73] | Internal capsule | Right Posterior |
| | | 284 | 4.04 | 0.012 | 0.16 | [66 101 83] | | Right Posterior |
| | | 2392 | -5.09 | <0.001 | 0.20 | [125 114 60] | External capsule | Left |
| | | 1767 | -4.69 | <0.001 | 0.18 | [92 91 83] | CC | Splenium |
| | | 294 | -3.99 | 0.009 | 0.15 | [87 124 98] | | Body |
| | | 1052 | -4.78 | <0.001 | 0.18 | [55 81 64] | Sagittal stratum | |
| | | 553 | -4.78 | <0.001 | 0.18 | [82 99 51] | | |
| | | 735 | -4.33 | <0.001 | 0.17 | [106 80 40] | Cerebellar peduncle | Middle |
| | | 339 | -3.88 | 0.002 | 0.15 | [74 79 41] | | Middle |
| | | 334 | -4.80 | 0.003 | 0.18 | [84 91 54] | | Right Superior |
| | | 727 | -5.03 | <0.001 | 0.19 | [128 103 100] | Longitudinal fasciculus | Left Superior |
| | | 658 | -5.03 | <0.001 | 0.19 | [58 113 107] | | Right Superior |
| | | 689 | -3.68 | <0.001 | 0.14 | [116 114 95] | Corona radiata | Left Superior |
| | | 554 | -4.72 | <0.001 | 0.18 | [63 100 70] | Fornix | Right |
| | | 413 | -4.62 | <0.001 | 0.18 | [96 109 51] | Cerebral peduncle | Left |
| | Fluid Composite | 1012 | 4.66 | <0.001 | 0.18 | [68 98 92] | Corona radiata | |
| | | 603 | 4.33 | <0.001 | 0.17 | [81 78 79] | | |
| | | 263 | 4.06 | 0.009 | 0.16 | [50 89 60] | Sagittal stratum | Right |
| | | 248 | 3.99 | 0.015 | 0.15 | [124 113 58] | Fornix | Left |

| NODDI metric | Cognitive test | Voxels | Peak t-stat | P-Value | Peak Cohen's d | Coordinates [X Y Z] | Region | Subregion |
|---|---|---|---|---|---|---|---|---|
| | | 4437 | -4.96 | <0.001 | 0.19 | [83 91 80] | CC | Splenium |
| | | 1053 | -4.67 | <0.001 | 0.18 | [129 95 100] | Longitudinal fasciculus | Left Superior |
| | | 850 | -3.94 | <0.001 | 0.15 | [113 75 34] | Cerebellar peduncle | Middle |
| | | 543 | -4.25 | <0.001 | 0.16 | [73 81 36] | | Middle |
| | | 527 | -4.75 | <0.001 | 0.18 | [81 101 32] | | Middle |
| | | 449 | -3.88 | <0.001 | 0.15 | [126 88 78] | Internal capsule | Left Retrolenticular |
| | Early Childhood Composite | 722 | 4.19 | <0.001 | 0.16 | [67 157 73] | Corona radiata | Right Anterior |
| | | 523 | 3.84 | <0.001 | 0.15 | [64 101 106] | | Right Superior |
| | | 474 | 3.68 | <0.001 | 0.14 | [105 164 74] | | Left Anterior |
| | | 405 | 4.22 | <0.001 | 0.16 | [111 131 99] | | Left Superior |
| | | 548 | 3.89 | <0.001 | 0.15 | [97 83 93] | CC | Splenium |
| | | 372 | 3.73 | <0.001 | 0.14 | [77 86 99] | | Splenium |
| | | 529 | 4.85 | <0.001 | 0.19 | [68 98 92] | Corona radiata | |
| | | 412 | 4.30 | <0.001 | 0.17 | [55 111 58] | Fornix | Right |
| | | 5096 | -4.47 | <0.001 | 0.17 | [88 124 98] | CC | Body |
| | | 1289 | -4.99 | <0.001 | 0.19 | [128 95 100] | Longitudinal fasciculus | Left Superior |
| | | 250 | -3.86 | 0.022 | 0.15 | [60 98 110] | | Right Superior |
| | | 1089 | -4.99 | <0.001 | 0.19 | [80 97 31] | Cerebellar peduncle | Middle |
| | | 609 | -4.02 | <0.001 | 0.15 | [113 75 34] | | Middle |
| | | 500 | -4.83 | <0.001 | 0.19 | [59 104 73] | Internal capsule | Right Retrolenticular |
| | | 377 | -4.09 | <0.001 | 0.16 | [99 104 46] | Corticospinal tract | Left |
| | | 368 | -4.00 | <0.001 | 0.15 | [83 105 50] | | Right |
| | | 268 | -3.53 | 0.012 | 0.14 | [57 120 59] | External capsule | Right |
| | | 262 | -3.46 | 0.015 | 0.13 | [123 118 67] | | Left |

Grey cells indicate a positive t-statistic and white cells a negative t-statistic. Significance was determined using a permutation cluster-wise $P < 0.050$ familywise error (FWE)-corrected threshold. Positive t-statistic values indicate a significantly positive regression between the diffusion signal and cognitive test, while negative t-statistic values indicate a significantly negative regression in the HCP control group (n = 679, ages 22–30). *Abbreviations: [X Y Z], coordinates of the voxels on the MNI 1 mm brain atlas; CC, corpus callosum; CCC, Cognitive Crystallized Composite; CS, cancer survivors; FWE, familywise error; HC, healthy controls; HCP, Human*

*Connectome Project; ISO, isotropic volume fraction; NDI, neurite density index; ODIp, primary orientation dispersion index; ODIs, secondary orientation dispersion index; ODIt, total orientation dispersion index; t, t-statistic; WM, white matter.*
